# Fine-tuning biosensor dynamic range based on rational design of cross-ribosome-binding sites in bacteria

**DOI:** 10.1101/2020.01.27.922302

**Authors:** Nana Ding, Shenghu Zhou, Zhenqi Yuan, Xiaojuan Zhang, Jing Chen, Yu Deng

## Abstract

Currently, predictive translation tuning of regulatory elements to the desired output of transcription factor based biosensors remains a challenge. The gene expression of a biosensor system must exhibit appropriate translation intensity, which is controlled by the ribosome-binding site (RBS), to achieve fine-tuning of its dynamic range (i.e., fold change in gene expression between the presence and absence of inducer) by adjusting the translation initiation rate of the transcription factor and reporter. However, existing genetically encoded biosensors generally suffer from unpredictable translation tuning of regulatory elements to dynamic range. Here, we elucidated the connections and partial mechanisms between RBS, translation initiation rate, protein folding and dynamic range, and presented a rational design platform that predictably tuned the dynamic range of biosensors based on deep learning of large datasets cross-RBSs (cRBSs). A library containing 24,000 semi-rationally designed cRBSs was constructed using DNA microarray, and was divided into five sub-libraries through fluorescence-activated cell sorting. To explore the relationship between cRBSs and dynamic range, we established a classification model with the cRBSs and average dynamic range of five sub-libraries to accurately predict the dynamic range of biosensors based on convolutional neural network in deep learning. Thus, this work provides a powerful platform to enable predictable translation tuning of RBS to the dynamic range of biosensors.

## INTRODUCTION

Biosensors have gained major attention in the field of biotechnology ^1^ especially for monitoring metabolite formation ^2, 3^. Genetically encoded biosensors derived from small-molecule inducer responsive transcription factors that produce fluorescence intensity proportional to the target metabolite concentration in the detection range have attracted substantial research attention ^3, 4^. However, the existing genetically encoded biosensors generally have the drawback of inappropriate dynamic range (i.e., fold change in gene expression between the presence and absence of inducer) ^5–9^. Dynamic range is an important indicator for fine-tuning biosensors, and a high dynamic range can help to distinguish the small difference in the inducer concentrations. The gene expression in biosensor systems driven by small molecule responsive transcription factors can achieve the desired output at appropriate translation initiation rates (TIR). One of the key elements to regulate the TIR is the ribosome-binding site (RBS), which tunes the dynamic range of the biosensor by adjusting the TIR of the transcription factor and reporter. However, the existing genetically encoded biosensors usually suffer from unpredictable translation tuning of regulatory elements to dynamic range. Many attempts have been made to tune the dynamic range of biosensors. For instance, Levin-Karp *et al*. used six RBSs ranging from strongest to weakest to achieve 20–200-fold dynamic range of protein expression ^10^. Wang *et al*. tuned the dynamic range of device input and output using five various-strength RBSs (RBS30–RBS34) from the Registry of Standard Biological Parts, and showed that RBS could be used as a linear amplifier to regulate protein expression levels ^11^. Although these methods might help to regulate the dynamic range of gene expression, the dynamic range of regulatory elements involved in gene expression could not been predicted. For example, if the RBS was changed, then obtaining the appropriate dynamic range of gene expression required time-consuming and laborious research.

Establishment of a predictable and robust method can quickly achieve translation tuning of the RBS to biosensor dynamic range. In a previous report, Salis *et al*. calculated the Gibbs free energy difference (ΔG_tot_) between the initiation and termination states of protein translation initiation based on a thermodynamic model, and presented RBS calculator for designing and synthesizing the RBSs of genes of interest, ensuring the rational control of protein expression levels ^12^. This significant contribution had accelerated the construction and optimization of complex genetic systems as well as promoted the development of synthetic biology. However, synthesis of the RBS through the calculation of free energy lacked experimental support. Therefore, rational design of the RBS by using a large amount of experimental data could make research on the RBS synthesis more robust. However, a large RBS database must rely on powerful analysis tools for better utilization of their application value, which can be solved by using mathematical models such as deep learning. Deep learning is an algorithm that uses artificial neural networks as a framework to characterize and learn databases. Deep learning models based on sequence levels have broad application prospects in the field of synthetic biology. For example, Chen *et al*. established Selene, a PyTorch-based deep learning library, which enables researchers to easily train the existing models to process biological problems of interest based on new databases and can be applied to any biological sequence data, including DNA, RNA, and protein sequences ^13^. Nielsen and Voigt used a deep learning based convolutional neural network (CNN) containing 42,364 plasmid DNA sequences datasets from Addgene to predict the lab-of-origin of a DNA sequence, and achieved 70% prediction accuracy and rapid analyses of DNA sequence information to guide the attribution process and understand the measures ^14^. While these studies provide a window for translation tuning of the RBS to biosensors dynamic range, the ability to design biosensors with reasonable dynamic ranges still remains a challenge ^15–17^.

In general, the RBS controls the translation initiation rate of a protein, thus affecting the protein expression level ^12^. Therefore, in the study of biosensors, the RBS tunes the dynamic range of biosensors by regulating the expression of reporter and regulatory protein. In the present study, the RBS design principles for *cdaR* and *sfgfp* in glucarate biosensors were established. Subsequently, a library containing 24,000 cross-RBSs (cRBSs, combining RBSs of *cdaR* and *sfgfp* in glucarate biosensors) was constructed by using DNA microarray, which was divided into five sub-libraries through fluorescence-activated cell sorting (FACS). Finally, a CNN on the cRBSs libraries was trained and a classification model between cRBSs and average dynamic range of each sub-library was developed and was termed CLM-RDR, which performed well in predicting biosensors dynamic range (**Fig. 1**). The CLM-RDR used large RBS data according to a semi-rational design to provide a knowledge base for precise adjustment of biosensors dynamic range, thus helping researchers to better characterize biosensors dynamic range by using RBS datasets. Given the availability of a large number of semi-rationally designed RBSs, the CLM-RDR classification model can be extended to other biosensors to fine-tune their dynamic ranges, thereby significantly simplifying the workload of the design–build–test–learn cycle for designing biosensors with moderate dynamic ranges in bacteria and accelerating intelligent fine-tuning of biosensor dynamic range.

**Fig. 1.**
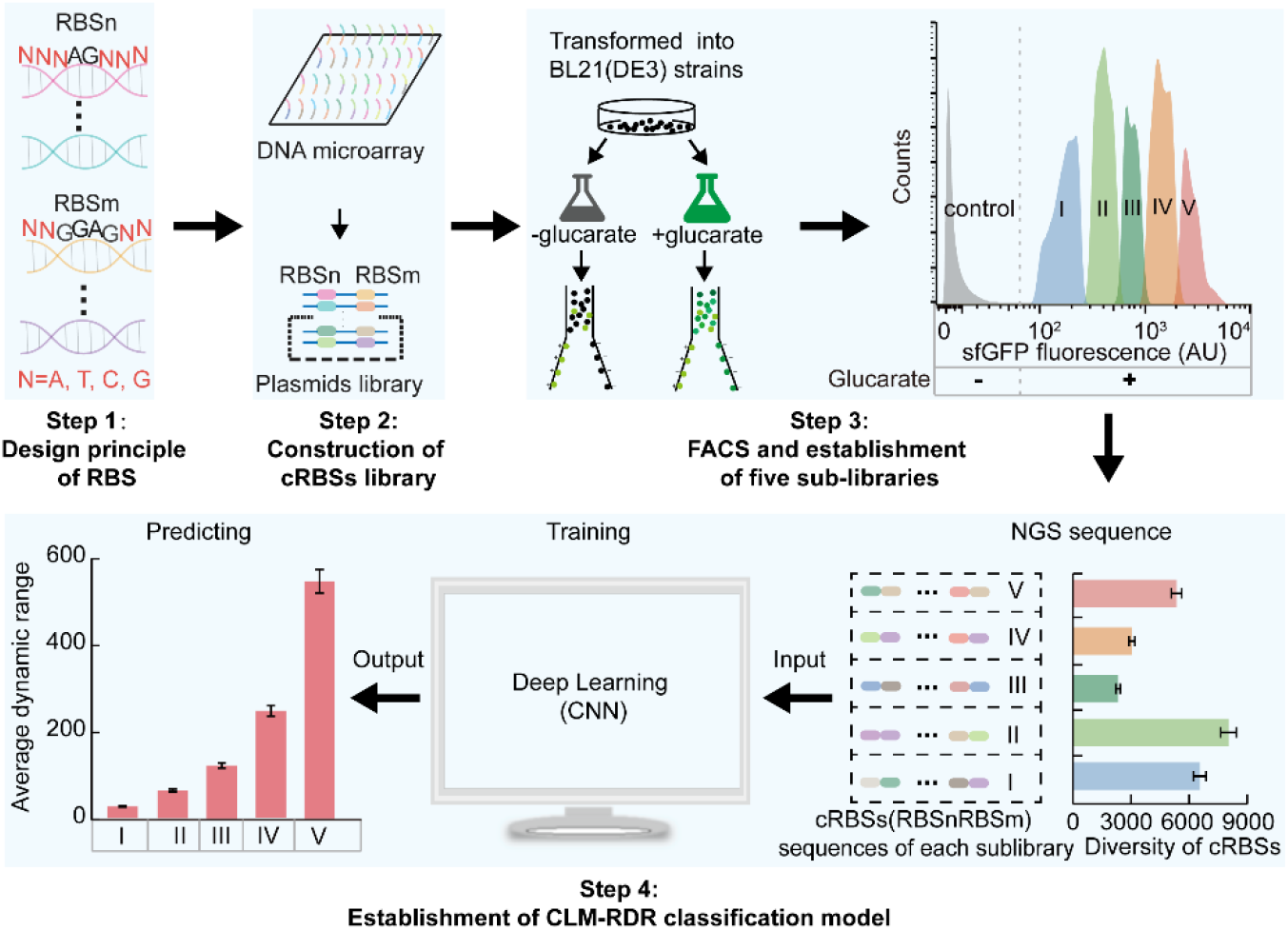
Workflow of CLM-RDR development.

First, the dynamic range of biosensors and the sequences of their related cRBSs were analyzed to establish an RBS design principle (**Step 1**). Based on this principle, a cRBSs library was designed and synthesized (**Step 2**) using DNA microarray. Subsequently, the library was divided into five sub-libraries (I–V) based on the fluorescence intensity of sfGFP measured by FACS (**Step 3**). Finally, to predict the dynamic range of biosensors with the given cRBSs, NGS and CNN model were employed to analyze the sequences of cRBSs in sub-libraries I–V and establish the CLM-RDR, respectively (**Step 4**). RBSn (NNNAGNNN), RBSs of *cdaR*; RBSm (NNGGAGNN), and RBSs of *sfgfp*; N = A, T, C, G.

## RESULTS

### RBS plays a crucial role in the regulation of biosensor dynamic range

Although recent advances in synthetic biology have shed light on the importance of fine-tuning of biosensor dynamic range in various fields, the ability to design biosensors with moderate dynamic ranges remains limited ^9, 18–20^. To investigate the key factors in biosensor dynamic range regulation, we used glucarate biosensor and explored its response strength by employing diverse concentrations of glucarate for induction (**Supplementary Fig. 1a, b**). Addition of 20 g/L glucarate biosensor presented the highest nine-fold dynamic range. However, the fluorescence intensity presented a downward trend when the glucarate concentration exceeded 20 g/L (**Supplementary Fig. 1b**). Similar observations have also been noted for other biosensors, such as *acuR*-based 3-hydroxypropionate biosensor ^3^, which also exhibited downward trend of fluorescence intensity when cerulenin concentration exceeded a certain threshold value. This phenomenon may be owing to the rapid translation and transcription of sfGFP, which not only cause metabolic burden (slow growth) (**Supplementary Fig. 1c**) to the living cells, but also affect the natural folding of sfGFP ^21^, thus resulting in low fluorescence intensity. Faure *et al*. indicated that the occurrence of misfolding proteins increases with the increasing translation speed ^22^. Thus, although the amount of expressed sfGFP increased (**Supplementary Fig. 1d**), the fluorescence intensity per protein molecule significantly decreased when glucarate concentration exceeded 20 g/L, owing to excessive misfolding. A similar trend was also observed for CdaR. Therefore, it can be assumed that the most critical challenge for fine-tuning the dynamic range of biosensors might be to balance the translation rate of regulator and reporter to simultaneously achieve the desired total fluorescence intensity with the highest fluorescence intensity per protein molecule (**Fig. 2a**). These findings suggested that RBS might probably be a key element affecting the dynamic range of biosensors.

**Fig. 2.**
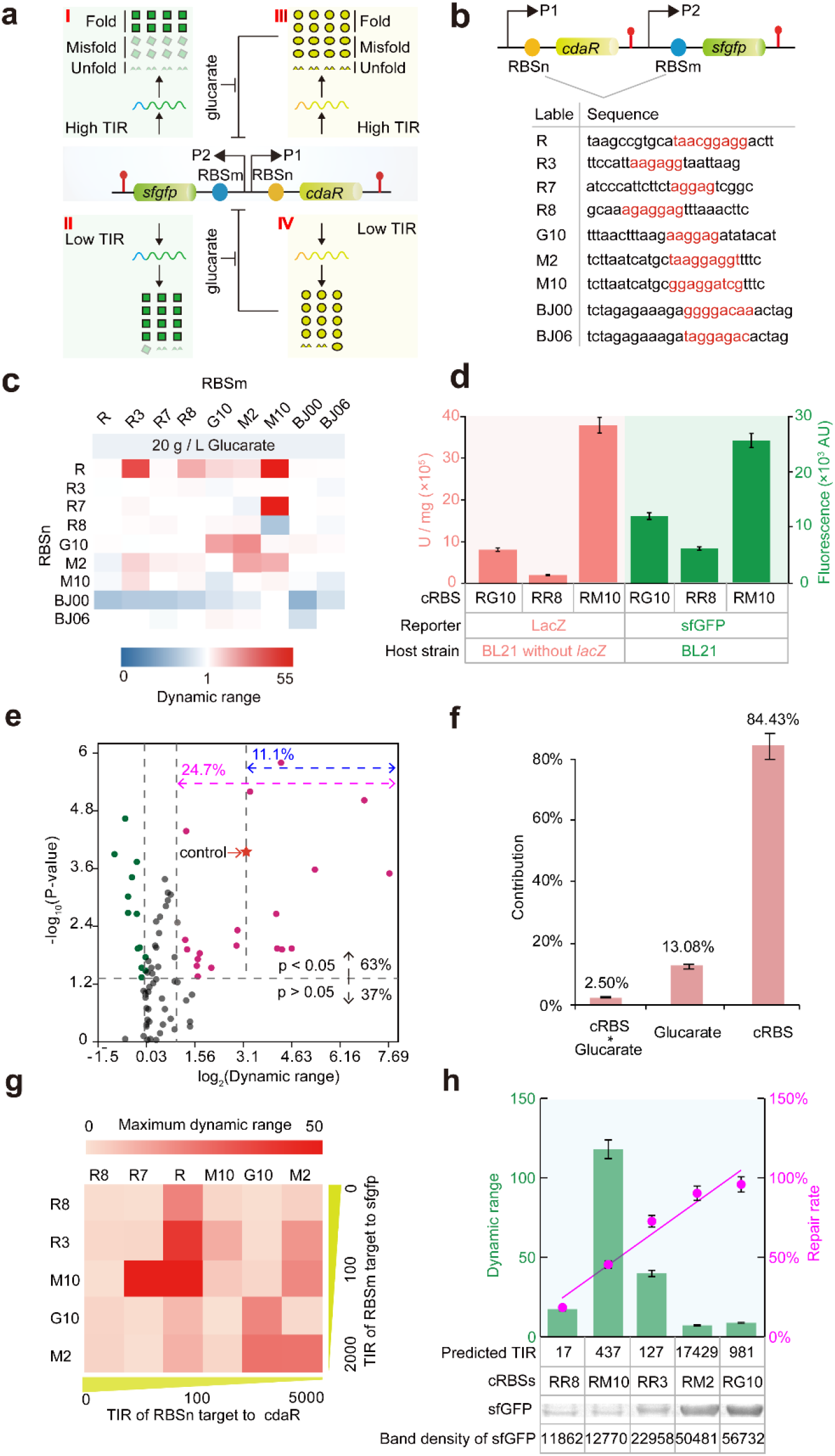
Effects of cRBSs on biosensor dynamic range.

(**a**) Hypothesis indicating that RBS affects protein folding. TIR coordinates both protein expression and folding state. (**b**) Nine RBS sequences derived from various libraries were obtained to replace the RBSs of glucarate biosensor.(**c**) The dynamic ranges of 81 cRBS glucarate biosensors induced with different concentrations of glucarate. cRBSs are defined as the RBS combination of *cdaR* (RBSn) and *sfgfp* (RBSm); for example, RM10 (R represents the RBSn of *cdaR*, M10 denotes the RBSm of *sfgfp*). (**d**) Comparison of LacZ enzyme activity and sfGFP expression intensity in three cRBS glucarate biosensors controlled by two reporter genes. Red column, LacZ enzyme activity; Green column, fluorescence intensity. (**e**) Volcano plot of cRBS datasets. The horizontal gray dashed line indicates a P-value of 0.05. The upper part (P < 0.05) represents the significant cRBS datasets. The vertical gray dashed line from left to right denotes the onefold, twofold, and nine-fold dynamic range. The pink and blue double arrows represent the significantly different cRBS datasets. Red star indicates the ninefold dynamic range of the control cRBS (RG10). (**f**) ANOVA for mean-normalized dynamic range from cRBSs and glucarate concentration datasets, with element- and junction-specific contributions to total dynamic range as noted (**Materials and Methods**). (**g**) Effect of TIR on the dynamic range of glucarate biosensor. Yellow triangle bars represent the increasing TIRs of RBSn and RBSm for *cdaR* and *sfgfp*, respectively. (**h**) Analysis of the dynamic range of biosensor and repair rate of sfGFP based on the distinct TIRs and sfGFP expression of RBSs. The correlation coefficient square (R^2^) of the fitted curve of the repair rate was 0.95. Band density was measured using ImageJ software; green columns represent the dynamic range of the biosensor; pink circles indicate the repair rate of sfGFP controlled by different cRBSs; repair rate is calculated as: (Flu (GroELS+) – Flu (GroELS–)) / Flu (GroELS+), where Flu denotes fluorescence intensity.

To investigate the correlation between RBS and biosensor dynamic range, nine RBSs covering a wide range of TIR from weak to strong were chosen for combinatorial replacement of the RBSs of *cdaR* and *sfgfp* (**Fig. 2b**). The nine RBSs selected were RBS (R) and G10RBS (G10) derived from the plasmid pJKR-H-*cdaR* ^4^; RBS3 (R3), RBS7 (R7), and RBS8 (R8) designed with an RBS calculator ^12^; MCD2 (M2) and MCD10 (M10) derived from the monocistronic design by Mutalik *et al*. ^*23*^; and BBa_J61100 (BJ00) and BBa_J61106 (BJ06) obtained from the Anderson RBS library. Finally, 81 cRBS glucarate biosensors were obtained and their response strength and dynamic range were significantly improved when induced with various concentrations of glucarate (**Fig. 2c, Supplementary Fig. 2a, b**). In the cRBSs of R7M10 and RM10, 205-fold and 118-fold dynamic ranges were observed, respectively, depending on glucarate concentration (20 g/L), which were higher than that of the control RG10 (9-fold), indicating that the RBS played a very important role in fine-tuning biosensor dynamic range.

To validate whether the effect of cRBSs on the biosensor dynamic range was independent of reporter genes, we selected three cRBS biosensors with distinct dynamic ranges (RG10, RR8, and RM10) to replace *sfgfp* with *lacZ*. By comparing LacZ enzyme activity and sfGFP expression intensity, we found that the three cRBSs showed the same expression intensity trend regardless of the reporter gene (*sfgfp* or *lacZ*) (**Fig. 2d**). This finding indicated that the cRBSs could consistently fine-tune the dynamic range of biosensor irrespective of the reporter. Subsequently, we analyzed the datasets with and without 20 g/L glucarate to assess the significance of differential expressions of genes with 81 cRBSs. We found that 63% of the 81 cRBSs were available for analysis (P < 0.05), and that 24.7% of the cRBSs showed significant differential expression (**Fig. 2e**). Moreover, 11.1% of the 81 cRBSs were significantly differentially expressed, when compared with the control (RG10) (**Fig. 2e**). To verify whether RBS was the most critical factor affecting the dynamic ranges of glucarate biosensors, we performed analysis of variance (ANOVA) on cRBSs and glucarate datasets (**Fig. 2f**). The results suggested that cRBSs and glucarate contributed 84% and 13% to biosensor fine-tuning, respectively. In addition, an interaction (2%) between the two factors was also noted (**Supplementary Table 1, see online methods**). These results indicated that the RBS is a key element for tuning the dynamic range of biosensors. However, it is still unclear on how the RBS fine-tunes the biosensor dynamic range.

### The RBS fine-tunes biosensor dynamic range by controlling protein translation and folding

To explore the relationship between TIR and dynamic range, total Gibbs free energy of the two variables, RBSn and RBSm, were respectively analyzed by using the RBS calculator ^12^ (**Supplementary Table 2**). Under the same RBSn, the optimal TIR of RBSm produced the highest biosensor dynamic range, and similar trend was also found for the TIR of RBSn under the same RBSm (**Fig. 2g, Supplementary Fig. 2c, d**), suggesting that the maximum dynamic range can be achieved at optimal TIR. However, TIR higher than the optimal TIR could cause low biosensor dynamic range, which could be due to the rapid expression of sfGFP resulting in misfolding or unfolding, thus affecting the natural folding of sfGFP ^**22, 24**^. Therefore, we hypothesized that the RBS could affect protein folding by regulating the TIR of protein.

To examine the relationship between dynamic range and protein folding, the reported wild-type chaperone ring complex, GroEL/S, which has the ability to assist in the folding of heterologous protein in *Escherichia coli*^25^, was used to verify the effect of the RBS on sfGFP folding. Five cRBSs (RR8, RM10, RR3, RM2, and RG10) with different TIRs were used to investigate the misfolding and repair of sfGFP. The fluorescence changes with and without GroEL/S were explored by flow cytometry upon addition of 20 g/L glucarate **(Fig. 2h)**. SDS-PAGE revealed that the increase in fluorescence intensity of each cRBS was not caused by different expression levels of sfGFP, but was caused by GroEL/S repairing misfolded or unfolded sfGFP to a natural folded state (**Supplementary Fig. 2e**). Furthermore, the repair rate, dynamic range, TIR, and sfGFP expression levels were calculated, which indicated that sfGFP expression was positively correlated with repair rate, while optimal TIR was more beneficial for achieving higher biosensor dynamic range **(Fig. 2h, Supplementary Fig. 2f–2h)**. This finding was consistent with our hypothesis, implying that strong RBSs have high TIR, which not only promotes the translation of sfGFP, but also results in high misfolding rate and repair rate. Although dynamic range is a comprehensive phenomenon indicating the amounts and folding state of sfGFP, it is difficult to establish a quantitative equation to define the relationship between the RBS, TIR, folding, and dynamic range, which severely hinders the development of rational design of biosensors.

### Semi-rational design of the RBS to fine-tune biosensor dynamic range

Owing to the lack of quantitative relation between the RBS, TIR, folding, and dynamic range, it is possible to simulate and predict the biosensor dynamic range by mathematical models. As an alternative method, deep learning could predict complex biological relationships with simple neural network models, thereby circumventing the steps to understand the complicated biological mechanisms and achieving the expected effects of simulation and prediction. To obtain large data to train CNN model, we first accomplished rational designing of the RBS and further tuned the dynamic range of the biosensor. On the basis of the 81 cRBSs datasets, the conserved sequences of the RBSs in *cdaR* and *sfgfp* were generated by using the online software WebLogo ^26^. The engineered RBSs could be divided into a consensus sequence defined as upstream and downstream of the Shine-Dalgarno (SD) sequence (RBSn: TAACCATGCATA-SDn-GACTT for *cdaR*; RBSm: TCTTAATCATG-SDm-GGTTTC for *sfgfp*) and an SD preference sequence (SDn: NNGGAGNN for *cdaR*; SDm: NNNGANNN for *sfgfp*; N = A, T, C, G) (**Fig. 3a, b**).

**Fig. 3.**
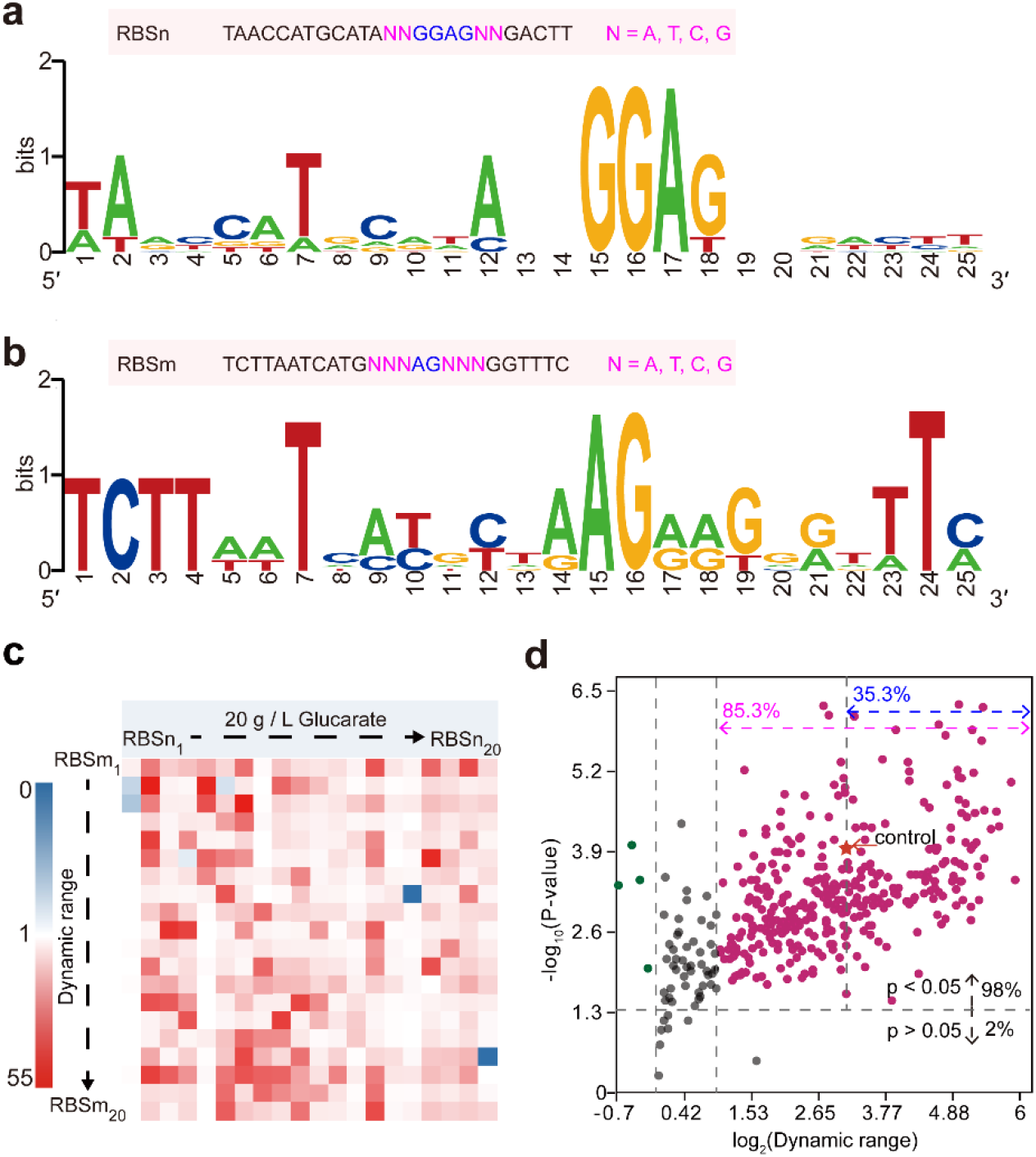
Semi-rational design of RBSs. The semi-rational design principle of (**a**) RBSn (RBSs of *cdaR)* and (**b**) RBSm (RBSs of *sfgfp*) was obtained based on the 81 cRBSs sequences using the online software WebLogo. (**c**) Biosensors dynamic ranges of 400 cRBSs, which were designed by the semi-rational design principle of RBSs, were calculated upon the addition of 20 g/L glucarate. (**d**) Volcano plot of 400 cRBSs datasets. The horizontal gray dashed line represents a P-value of 0.05. The upper part (P < 0.05) denotes the significant cRBS datasets. The vertical gray dashed line from left to right indicates onefold, twofold, and ninefold dynamic range. The pink and blue double arrows show the significantly different cRBS datasets. Red star represents the ninefold dynamic range of the control cRBS (RG10).

To evaluate the reliability of this design principle of RBSs, we randomly constructed 400 cRBSs (20 × 20 RBSs, 20 RBSs of *cdaR* and *sfgfp)* (**Supplementary Table 3**). The fluorescence intensity and dynamic range of the 400 cRBSs biosensors with glucarate inducer showed a significant improvement, when compared with those without the inducer (**Supplementary Fig. 3**). In addition, the cRBSs biosensors presented an improved dynamic range upon addition of 20 g/L glucarate, when compared with the control (**Fig. 3c**). These findings implied that semi-rational design of cRBSs was more reliable and robust in improving the biosensor dynamic range. We further analyzed the datasets with and without glucarate to assess the differential expression of sfGFP, and found that up to 98% of the 400 cRBSs were available for analysis (P < 0.05) and 85.3% of the cRBSs showed significant differential expression (**Fig. 3d**). In particular, 35.3% of the 400 cRBSs presented significant differential expression, when compared with the control (RG10) (**Fig. 3d**). These results indicated that the semi-rational design of cRBSs considerably contributed to the improvement of biosensor dynamic range.

### Establishment of CLM-RDR for precise prediction of biosensor dynamic range

To further extend the dataset for CNN model training, we constructed a much larger cRBS library through the RBS semi-rational design approach, and generated 100 RBSs for *cdaR* and 120 RBSs (**Supplementary Table 3**) for *sfgfp* (**Fig. 3a, b**). Then, a combinatorial library of 12,000 cRBSs as oligonucleotides was developed with DNA microarray (**see online methods**). To verify the homogeneity of the 12,000 cRBSs, next-generation sequencing (NGS) was performed. The coverage of the 12,000 cRBSs was 100%, and the 10-fold variation reached a quality control value of 99.92% (**Supplementary Fig. 4a, Supplementary Data 1, Accession No. SRR9301216**). This cRBS library was used in the following pooled screening experiment to characterize the dynamic range of the glucarate biosensor.

The 12,000 cRBS plasmid library was transformed into *Escherichia coli* (*E. coli*) BL21 (DE3) cells, which were cultured for 8 h in Luria–Bertani (LB) medium supplemented with 0 or 20 g/L glucarate. Then, by using FACS, we divided the cells induced with 20 g/L glucarate into five non-adjacent sub-libraries I–V according to the expression intensity of sfGFP, and compared them with the control without glucarate induction (**Fig. 4a**). Subsequently, the average single cell fluorescence intensity and average dynamic range of the sub-library I–V and control were calculated, and a 26-fold, 63-fold, 121-fold, 246-fold, and 545-fold average dynamic range were accomplished for the sub-libraries I–V, respectively (**Fig. 4b**). These results further demonstrated that the cRBS semi-rational design approach was highly effective in tuning the dynamic range of the glucarate biosensor, and helped to establish a high-quality element library in synthetic biology and construct an approach for designing complex genetic circuits to fine-tune gene expression ^27–29^.

**Fig. 4.**
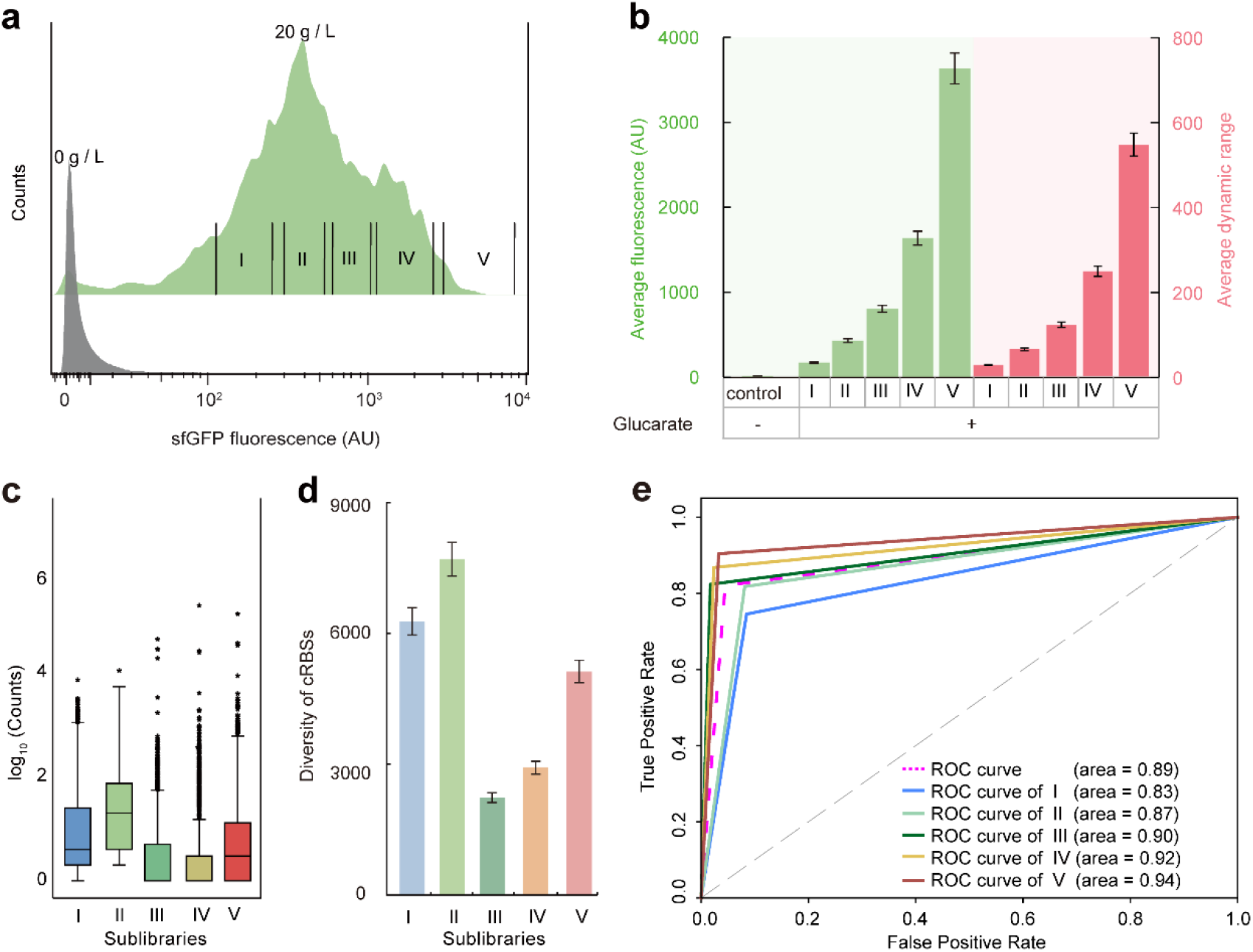
Accurate prediction of the dynamic range of glucarate biosensor from cRBS sequences by deep learning model. (**a**) A larger cRBSs library was formed than the original libraries. Division of cells induced with 20 g/L glucarate into five non-adjacent sub-libraries (I–V), which were compared with the control (0 g/L glucarate) based on the expression intensity of sfGFP measured by FACS. (**b**) Analysis of average fluorescence intensity (green column) and average dynamic range (red column) of each sub-library and control. (**c**) The counts of each cRBS of the five sub-libraries were obtained by NGS. (**d**) Diversity of cRBSs of five sub-libraries. (**e**) Establishment of CLM-RDR based on 24,000 cRBS sequences. Receiver operating characteristic (ROC) curves for cRBSs of sub-libraries I–V (solid lines of various colors) and total library (pink dotted line). Biosensor dynamic ranges with five test-positive samples were used to classify.

To determine the cRBS sequences of the glucarate biosensors in each sub-library, we first obtained the assorted biosensor plasmids of the five sub-libraries. Then, the mixed PCR products of the five modified sub-libraries were linked with five barcodes and sequenced by NGS ^30^ (**Accession No. SRR9301175**; **see online methods**). Box plots showed the distribution of each cRBS count of five sub-libraries, and separate points indicated that the cRBS numbers ranged from 10 to 10^5^ (**Fig. 4c, Supplementary Data 2**). In addition, the diversity of cRBSs in each sub-library was analyzed, and there were 6219, 7630, 2214, 2892, and 5079 cRBSs in sub-libraries I–V, respectively (**Fig. 4d**). Besides, more than 12,000 cRBSs were found, possibly because of mutations introduced into the sequence through bacterial evolution during cultivation. Although the mutation rates of the consensus sequences of RBSn and RBSm in the five sub-libraries were 0.15, 0.19, 0.06, 0.09, and 0.15, respectively, and they did not affect subsequent model development (**Supplementary Data 2**). Therefore, to ensure data integrity, the sequenced 24,000 cRBSs were used as the data sources for further data processing.

Although the cRBSs sequences of each sub-library were obtained, it was extremely crucial to determine the functional relationships between the cRBSs sequences and average dynamic range of glucarate biosensor. Functional relationships could help to quickly analyze the dynamic range of a corresponding cRBS biosensor, which could reduce the burden of the design–build–test–learn cycle. Therefore, CNNs of deep learning was chosen to establish a classification model between cRBSs and the average dynamic range of each sub-library (CLM-RDR). The cRBSs and average dynamic range of sub-libraries I–V were the input and output of CLM-RDR, respectively. First, 85% of the cRBSs in each sub-library were randomly selected as datasets to train the CNN model (**Supplementary Fig. 5**). Next, we evaluated how well CLM-RDR predicted the average dynamic range of the glucarate biosensor from the remaining 15% of cRBSs sequences in each sub-library (**Fig. 4e**). The results indicated that CLM-RDR predicted the dynamic range of the glucarate biosensor with high accuracy, yielding an area under the curve (AUC) of 0.83, 0.87, 0.90, 0.92, and 0.94 for sub-libraries I–V, respectively, and an average AUC of 0.89. Moreover, CLM-RDR performed better in predicting sub-libraries with high dynamic range, when compared with that with low dynamic range, implying that cRBSs in the high dynamic range could more easily achieve fine tuning of the biosensor dynamic range.

### Applications of the CLM-RDR to other biosensors

The CLM-RDR is expected to tune the dynamic range of different biosensors. Therefore, to further evaluate the performance of the CLM-RDR, we randomly selected 16 cRBSs to modify the glucarate biosensor, glycolate biosensor, and arabinose biosensor (**see online methods**). We first predicted the average dynamic range of 16 cRBSs by using CLM-RDR and then performed an experiment to detect the dynamic ranges of the biosensors via FACS (**Supplementary Fig. 6**). By analyzing the predicted and experimentally observed dynamic ranges, CLM-RDR was found to have good predictive performance for three biosensors. Predicted accuracy rates of 62.5% (**Fig. 5a**), 62.5% (**Fig. 5b**), and 68.75% (**Fig. 5c**) were obtained for glucarate, arabinose (**Fig. 5d**), and glycolate (**Fig. 5e**) biosensors, respectively. These results indicated that the CLM-RDR had a certain degree of universality in predicting the dynamic ranges of biosensors. The CLM-RDR can probably be further improved by providing additional training datasets.

**Fig. 5.**
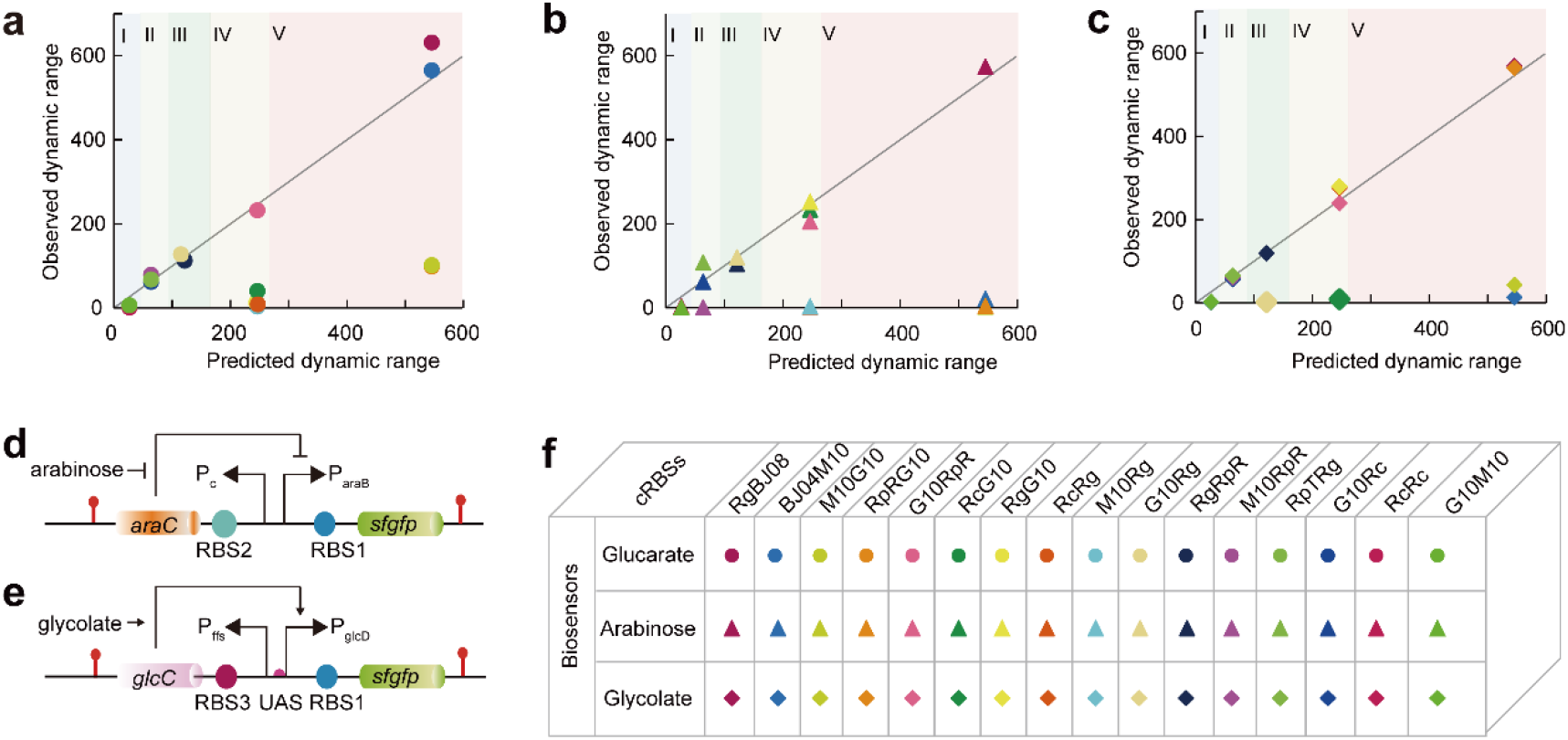
CLM-RDR verification for three genetically encoded biosensors. Sixteen cRBSs were randomly selected for biosensor modification and comparison of the observed and predicted dynamic ranges. The CLM-RDR performed well in predicting the dynamic ranges of (**a**) glucarate biosensor, (**b**) arabinose biosensor, and (**c**) glycolate biosensor. I–V represent the five sub-libraries of cRBSs. The black diagonal denotes y = x. (**d)** Structure of P_araB_-based arabinose sensor. Pc represents the constitutive promoter that controls transcription of the regulatory protein AraC. P_araB_ is an inducible promoter containing the AraC-binding DNA sequence. Blunt-end arrows denote repression. (**e**) Structure of P_glcD_-based glycolate sensor. P_ffs_ ^31^ indicates the constitutive promoter that controls transcription of the regulatory protein GlcC. P_glcD_ is a constitutive promoter that controls the transcription of the reporter sfGFP. In the absence of glycolate, GlcC remained as a non-functional regulatory protein, whereas in the presence of glycolate, the regulatory protein GlcC and glycolate bound to the activator GlcC-glycolate, which in turn bound to the upstream activation site (UAS) of the promoter PglcD, thus enhancing transcription and expression of *sfgfp*. Pointed arrows indicate activation. (**f**) Detailed illustration of 16 cRBSs and three biosensors. Solid circle: glucarate biosensor; solid triangle: arabinose biosensor; solid diamond: glycolate biosensor.

### Software package

To encourage experimental biologists to use CLM-RDR, we uploaded the model to GitHub, which converted an RBS sequence directly into biosensor dynamic range. The code for predicting biosensor dynamic range can be found at https://github.com/YuDengLAB/CLM-RDR.

## DISCUSSION

Genetically encoded biosensors derived from transcription factors responding to small-molecule inducers are receiving increasing research attention ^3^. The currently available genetically encoded biosensors usually have the major problem of inappropriate dynamic range ^6, 8^. Although many valuable works, such as promoter modification studies, have attempted to tune the dynamic range of biosensors, universality may be difficult to achieve owing to small datasets and insufficient analysis tools. Therefore, fine-tuning of the biosensor dynamic range remains a huge challenge ^5, 17^. In general, RBS controls the translation initiation rate ^12, 23^ of regulatory proteins and reporters, which can control the dynamic range of biosensors. Previous reports had indicated that the dynamic ranges of device input or output were not well tuned by replacing the RBS ^10^, mainly because the RBS design was not sufficiently rational and the RBS datasets were limited. Therefore, to fine-tune the dynamic range of biosensors, in the present study, we established the design principle of the RBS in biosensors through ANOVA and online WebLogo processing. Accordingly, 12,000 cRBSs were semi-rationally designed based on the design principle, and five average dynamic ranges were calculated by dividing the cRBSs into five sub-libraries using FACS. Most importantly, we developed CLM-RDR, a classification model between cRBSs and average dynamic range of five sub-libraries. The CLM-RDR showed accurately predictive performance and was able to quickly determine the average dynamic range of a biosensor corresponding to a cRBS. In addition, the CLM-RDR also had good predictive ability toward glycolate and arabinose biosensors, thus indicating that this model can be extended to other biosensors. Besides, the developed model significantly simplified the workload of the design–build–test–learn cycle of fine-tuned biosensor dynamic range in bacteria and accelerated intelligent fine-tuning of biosensor dynamic range.

RBSs play a role in fine-tuning genetic components and determining the TIR of proteins ^12, 23^. Proteins usually present tight and loose structures. The mRNA structure affects the translation rate of a protein, and fast translation prevents the formation of compact structures, which affects protein folding ^22^. Thus, we hypothesized that the RBS might also affect the conformations of proteins by controlling TIR, thereby achieving fine-tuning of gene expression. To further explore the relationship between TIR, protein folding, and biosensor dynamic range, a wild-type chaperone GroEL/S, which could assist in the folding of recombinant sfGFP in *E. coli*, was combined with a set of constructed biosensors ^25^. Although there was no one-to-one relationship between the protein expression level and predicted TIR, a positive correlation trend was noted. In other words, when compared with low TIR, high TIR not only increased the protein expression, but also produced more misfolded proteins, which in turn resulted in a higher repair rate of sfGFP by GroEL/S (**Fig. 2a, h**). Therefore, appropriate protein expression level and protein folding state achieved the optimal biosensor dynamic range, thus further implying that RBS is one of the key factors affecting the dynamic range of biosensors.

Sequence-based deep learning models had been reported to show good predictive performance for biological phenotypes ^13, 32, 33^. Deep learning models can accurately establish the correspondence between genotypes and phenotypes through large datasets, thus making investigations more universal. The present study found that one of the key factors affecting the dynamic range of biosensors was RBS. However, the mechanism of the RBS tuning the dynamic range of biosensors was complex (**Fig. 2a**), not only requiring exploration of the mechanism of RBS tuning translation and folding of regulators and reporter, but also examination of the binding mechanism of regulators and operator sites and further investigation of the effects on downstream reporter transcription. Therefore, analysis of these mechanisms using current technology is a huge challenge. However, deep learning models do not require understanding of specific mechanisms to establish the relationship between RBS and biosensor dynamic range, and can be extended to other biosensors research. Hence, to develop a universal tool to fine-tune the dynamic range of biosensors, we developed CLM-RDR, a classification model based on deep learning between cRBSs and average dynamic range. The CLM-RDR showed good prediction performance for the dynamic range of the biosensor using only less than 24,000 cRBSs datasets. More importantly, it could be extended to other biosensors, achieving the same prediction effects, implying that CLM-RDR has certain universality in predicting the dynamic range of biosensors. It should be noted that the present study only examined the effect of the RBS on biosensor dynamic range. The results of this study, along with further research on promoters, plasmid copy numbers, and regulatory protein evolution, could propel fine-tuning of the dynamic range of biosensors into the era of intelligence.

## ONLINE METHODS

### Strains and culture conditions

All strains used in this study are listed in **Supplementary Table 4**. *E. coli* JM109 and *E. coli* BL21 (DE3) cells were used for plasmid cloning and protein expression, respectively. M9 minimal medium, consisting of Na_2_HPO_4_ (6.78 g/L), KH_2_PO_4_ (3.0 g/L), NaCl (0.5 g/L), MgSO_4_·7H_2_O (0.5 g/L), CaCl_2_ (0.011 g/L), NH_4_Cl (1.0 g/L), and glucose (5 g/L), was used for fluorescence intensity assessment. The final concentrations of ampicillin, kanamycin, and spectinomycin employed in this study were 100, 50, and 50 μg/mL, respectively. The final concentration of isopropyl β-D-thiogalactoside was 1 mM.

### Plasmid construction

All plasmids and primers used in this study are listed in **Supplementary Tables 4 and 5**, respectively. The pJKR-H-*cdaR* plasmid for glucarate biosensor was purchased from Addgene (#62557). In addition to RBS and g10RBS, we selected seven RBSs: RBS3, RBS7, RBS8, MCD2, MCD10, BBa_J61100 and BBa_J61106 (**Supplementary Table 2)**. The primer design was based on the different RBS sequences, and the pJKR-H-*cdaR* plasmid was used as the template for plasmid PCR. Plasmids pJKR-H-RBSs-*cdaR*-RBSs (RBSs are represented as R, R3, R7, R8, G10, M2, M10, BJ00, or BJ06), pJKR-H-RBSn_81_-*cdaR*-RBSm_56_, pJKR-H-RBSn_81_-*cdaR*-RBSm_97_, and pJKR-H-RBSn_81_-*cdaR*-RBSm_117_ were constructed through DpnI digestion, and the digestion products were introduced into *E. coli* JM109 cells for screening by colony PCR and Sanger sequencing. The plasmids pJKR-H-R-*cdaR*-G10-*lacZ*-his, pJKR-H-R-*cdaR*-M10-*lacZ*-his, pJKR-H-R-*cdaR*-R8-*lacZ*-his, NGS-RBSn-RBSm-I, NGS-RBSn-RBSm-II, NGS-RBSn-RBSm-III, NGS-RBSn-RBSm-IV, NGS-RBSn-RBSm-V, and pRSF-*groEL-groES* were constructed using with Gibson assembly ^34^. The plasmid pHS-AVC-LW1125 was synthesized by Beijing Syngentech Co., Ltd in china. through DNA microarray technique.

Plasmids containing the glycolate biosensor pUC-*glcC-ffs* and arabinose biosensor pUC-*araC* were constructed through Gibson assembly methods. In both of the biosensors, the *rrnB* strong terminator, antibiotic resistance gene, and origin of replication were derived from the glucarate biosensor (pJKR-H-*cdaR*) ^4^. All the sequences of transcriptional regulators and their promoters are provided in **Supplementary Table 6**. To evaluate the general performance of the CLM-RDR, we randomly selected eight RBSs to engineer three biosensors using plasmid PCR method: RBS_*cdaR*_ (R_c_) and g10RBS derived from the glucarate biosensor; BBa_J61104 (BJ04) and BBa_J61108 (BJ08) obtained from the Anderson RBS library; MCD10 generated from the monocistronic design by Mutalik, *et al;* RBS*glcC* (R_g_) obtained from the glycolate biosensor; and RBS_pRSF_ (R_pR_) and RBS_pTrc99a_ (R_pT_) derived from plasmids pRSF and pTrc99a, respectively (**Supplementary Table 6)**. The plasmid construction methods for each biosensor had been described earlier, and the concentrations of the inducers, glycolate and arabinose, were 70 and 20 mM, respectively (**Supplementary Fig.6)**.

### ANOVA model for cRBSs:glucarate combinatorial datasets

To understand the contribution and interaction between cRBSs and glucarate in the precise regulation of biosensors, we performed ANOVA ^23^ on the following linear model, using fluorescence data from sfGFP ^35^

Fluorescence_*ijk*_ = μ + C_*i*_ + G_*j*_ + (C:G)_*ij*_ + ε_*ijk*_

> for *i* = (1–81); *j* = (1–12)

where Fluorescence_*ijk*_ is the fluorescent output signal measured from the translation element, C_*i*_, and induced substrate glucarate, G_*J*_: (C:G)_*ij*_ represents any interaction between the *i*th translational element and *j*th concentration of glucarate; μ is the overall average signal; and ε_*ijk*_ is the error term for each C:G combination. The analysis output is presented in **Supplementary Table 1**.

### β-Galactose activity assays

The process of gene deletion in *E. coli* BL21 (DE3) cells was performed as described by Jiang *et al* ^36^. The sgRNA of *lacZ* is shown in **Supplementary Table 6**. An appropriate amount of fermentation broth was centrifuged at 8000 × *g* for 10 min at 4 °C, the supernatant was discarded, and the cells were collected. The cells were washed twice with cold lysis buffer (Tris–HCl; 0.01 M, pH 7.5). Then, the cells were resuspended in 2.5 mL of 0.01 mol/L Tris–HCl buffer (pH 7.5), and glass beads ^37^ and 50 μL of PMSF stock solution were added to the cell culture. The cell culture was oscillated six times at high speed for 15 s each and placed on ice intermittently. Subsequently, 2.5 mL of Tris–HCl buffer were added to the culture, and the supernatant collected after centrifugation at 8000 × *g* for 15 min at 4 °C was the crude enzyme solution. Next, 1 mM o-nitrophenyl-β-D-galactopyranoside (oNPG) solution was prepared with 50 mM oNPG. Approximately 10 μL of the diluted crude enzyme solution and 20 μL of the oNPG solution were added to 70 μL of Z-buffer (16.1 g/L Na_2_HPO_4_·7H_2_O, 5.5 g/L NaH_2_PO_4_·H_2_O, 0.75 g/L KCl, 0.246 g/L MgSO_4_,7H_2_O, and 2.7 mL β-mercaptoethanol; pH 7.0, stored at 4 °C) for 10 min at 30 °C. Then, 120 μL of 1 mol/L pre-cooled Na_2_CO_3_ were immediately added to stop the reaction and develop color. Finally, the absorbance was measured with a spectrophotometer at a wavelength of 420 nm. One unit of enzyme activity was defined as the amount of enzyme catalyzing the production of 1 μmol o-nitrophenol (oNP) per minute ^38, 39^.

Bovine serum albumin (BSA) was dissolved in Z-buffer at different dilutions (0.0–0.2 mg/mL BSA), and standard curves were generated. Crude enzyme (20 μL) was added to 200 μL of Bradford reagent, mixed, and its absorbance was determined at a wavelength of 595 nm. The crude enzyme concentration was calculated with a standard curve. The formula for calculating the enzyme activity was as follows. U/mg protein = OD420 × 1.7/(0.0045 × protein content × crude enzyme volume × time), where OD420 is the optical density of the product oNP at 420 nm, coefficient 1.7 is the corrected value of the reaction volume, coefficient 0.0045 is the optical density (OD) of 1 mM oNP solution, protein content is expressed in mg/mL, crude enzyme volume is expressed in mL, and time is shown in min.

### Fluorescence assays

The cells were grown overnight to saturation before being diluted into fresh LB medium at a ratio of 1:100 and incubated at 250 rpm and 37 °C. After 3 h, 100 μL of log-phase cells were transferred to 96-well plates and stock inducers were respectively added to achieve the desired induction concentrations. Different concentrations of glucarate, glycolate, and arabinose were obtained by diluting 100 g/L glucarate, 1 M glycolate, and 1 M arabinose mother liquor in 96-well plates. Before measurements, the cultures were diluted into 0.01 M phosphate buffered saline (PBS; pH 7.4) to ensure that the OD600 value was about 0.5. Measurements were performed using a Biotek HT plate reader (Winooski, VT, USA) under excitation wavelength of 485/20 nm and emission wavelength of 528/20 nm at 37 °C and rapid shaking. Fluorescence intensity was measured in arbitrary units (AU), and the OD was determined by absorbance. For a given measurement, normalized fluorescence was determined by dividing the fluorescence by OD. The ratio of fluorescence to absorbance at 600 nm was used to compensate for the changes in cell density over time and between experiments (AU/OD).

*E. coli* BL21 (DE3) cells containing the plasmid libraries were cultured to saturation, and then incubated at a concentration of 1% into 250-mL flasks containing LB medium at 250 rpm and 37 °C. After 2 h, inducers were added to the desired final concentration, and incubation was resumed for 12 h. The induced cultures were diluted into cold PBS and kept on ice until evaluation with a BD FACS AriaII cell sorter (Becton Dickinson) ^24^. At least 100,000 events were captured for each sample. BD FACSDiva software was used to divide the gate for sfGFP ^35^ (bandpass filter, 530/30 nm; blue laser, 488 nm).

### Construction of the RBS library and NGS analysis

In total, 12,000 cRBS sequences were synthesized using DNA microarray, amplified by PCR, and were cloned into a glucarate biosensor plasmid backbone (pHS-BVC-LW274 and pHS-BVC-LW276) via two-step Golden Gate assembly ^34^ (completed by Synbiotic Gene Company) to obtain the glucarate biosensor plasmid library. Next, the plasmid library was transformed into *E. coli* BL21 (DE3) cells, which were cultured for 8 h in LB medium with or without 20 g/L glucarate supplementation. Then, the cells induced with 20 g/L glucarate were divided into five non-adjacent sub-libraries (I–V), which were compared with the positive control without glucarate induction according to the fluorescence intensity of sfGFP by FACS. To ensure the reliability of fluorescence intensity, cell adhesion was removed by executing FSC-A/FSC-H and SSC-A/SSC-H operation. Finally, the cells from each sub-library were obtained. Although the distance between the two RBSs in the glucarate biosensor was 2208 bp, NGS was able to measure only up to 250 bp; therefore, Gibson assembly ^34^ was used to modify the plasmids of the five sub-libraries. The modified sub-libraries contained 134 bp between two RBSs (**Supplementary Fig. 4b**), and the mixed PCR products of the five modified sub-libraries were linked with five barcodes and sequenced by NGS. Finally, the cRBS sequences and sequence abundance of the five sub-libraries were determined.

### Deep learning

First, 24,000 cRBS sequences were combined to create datasets for subsequent deep learning. Then, the fluorescence intensity was divided into five levels for evaluating the biosensor corresponding to the RBS. To classify the RBS sequences, one-hot coding was initially employed. A neural network model ^32, 33^ consisting of three convolutional layers and three full connection layers was proposed to accurately classify the RBS sequences. The convolutional layers comprised stride 1 and the pooling layers were non-overlapping. The convolution layer included two functions: feature extraction and feature mapping. On the one hand, the input of each neuron was connected to the local receptive field of the previous layer, and the local features were extracted. After the local features were extracted, the positional relationships between them and other features were also determined. On the other hand, each computing layer of the network was composed of multiple feature maps, each feature maps into a plane, and all the neurons on the plane exhibited the same weight. The feature map used the ReLU function with a small kernel of the influence function as the activation function of the convolution network, so that it had an invariance of displacement.

### Software and graphics generation

Deep learning was performed with SciPy (1.0.0), NumPy (1.14.0), and TensorFlow (1.9.0) Python packages.

## AUTHOR CONTRIBUTIONS

N.D. and Y.D. conceived the study; N.D., X.Z., and Y.D. designed the research; N.D. performed the experiments; N.D., Z.Y., X.Z., S.Z., J.C., and Y.D. analyzed the data; and N.D., Z.Y., S.Z., and Y.D. wrote the manuscript.

## DATA AND MATERIALS AVAILABILITY

Raw data of NGS for DNA microarray and cRBSs of five sub-libraries have been deposited to the NCBI Short Read Archive, with Accession No. BioProject: SRR9301216 (https://dataview.ncbi.nlm.nih.gov/object/SRR9301216) and SRR9301175 (https://dataview.ncbi.nlm.nih.gov/object/SRR9301175), respectively. The code used to predict biosensor dynamic range can be found at https://github.com/YuDengLAB/CLM-RDR.

## ACKNOWLEDGMENTS

The authors thank Professor Chong Zhang of Tsinghua University for the valuable discussions. This work was supported by the National Natural Science Foundation of China (21877053), National First-class Discipline Program of Light Industry Technology and Engineering (LITE2018-24), Top-Notch Academic Programs Project of Jiangsu Higher Education Institutions (TAPP), Fundamental Research Funds for the Central Universities (JUSRP51705A, JUSRP11964), and Open Project Program of China-Canada Joint Lab of Food Nutrition and Health, Beijing Technology and Business University (BTBU) and Jiangsu Province Science Foundation for Youths (BK20150159).

## REFERENCES AND NOTES

1. Prindle, A. et al. A sensing array of radically coupled genetic ‘biopixels’. Nature 481, 39 (2012).

2. Eggeling, L., Bott, M. & Marienhagen, J. Novel screening methods—biosensors. Current opinion in biotechnology 35, 30–36 (2015).

3. Rogers, J.K. & Church, G.M. Genetically encoded sensors enable real-time observation of metabolite production. Proc Natl Acad Sci U S A 113, 2388 (2016).

4. Rogers, J.K. et al. Synthetic biosensors for precise gene control and real-time monitoring of metabolites. Nucleic Acids Research 43, 7648–7660 (2015).

5. Zhang, F., Carothers, J.M. & Keasling, J.D. Design of a dynamic sensor-regulator system for production of chemicals and fuels derived from fatty acids. Nature biotechnology 30, 354 (2012).

6. Nguyen, N.H., Kim, J.-R. & Park, S. Application of Transcription Factor-based 3-Hydroxypropionic Acid Biosensor. Biotechnology and Bioprocess Engineering 23, 564–572 (2018).

7. Skjoedt, M.L. et al. Engineering prokaryotic transcriptional activators as metabolite biosensors in yeast. Nature chemical biology 12, 951 (2016).

8. Cheng, F., Tang, X.L. & Kardashliev, T. Transcription factor-based biosensors in high-throughput screening: advances and applications. Biotechnology journal 13, 1700648 (2018).

9. Kasey, C., Zerrad, M., Li, Y., Cropp, T.A. & Williams, G.J. Development of transcription factor-based designer macrolide biosensors for metabolic engineering and synthetic biology. Acs Synthetic Biology 7, acssynbio.7b00287 (2017).

10. Levin-Karp, A. et al. Quantifying translational coupling in E. coli synthetic operons using RBS modulation and fluorescent reporters. ACS synthetic biology 2, 327–336 (2013).

11. Wang, B., Kitney, R.I., Joly, N. & Buck, M. Engineering modular and orthogonal genetic logic gates for robust digital-like synthetic biology. Nature communications 2, 508 (2011).

12. Salis, H.M., Mirsky, E.A. & Voigt, C.A. Automated design of synthetic ribosome binding sites to control protein expression. Nature biotechnology 27, 946 (2009).

13. Chen, K.M., Cofer, E.M., Zhou, J. & Troyanskaya, O.G. Selene: a PyTorch-based deep learning library for sequence data. Nature methods 16, 315 (2019).

14. Nielsen, A.A. & Voigt, C.A. Deep learning to predict the lab-of-origin of engineered DNA. Nature communications 9, 3135 (2018).

15. Westbrook, A.M. & Lucks, J.B. Achieving large dynamic range control of gene expression with a compact RNA transcription–translation regulator. Nucleic Acids Research 45, 5614–5624 (2017).

16. Doong, S.J., Gupta, A. & Klj, P. Layered dynamic regulation for improving metabolic pathway productivity in Escherichia coli. Proceedings of the National Academy of Sciences of the United States of America 115, 2964 (2018).

17. Fuzhong, Z. & Jay, K. Biosensors and their applications in microbial metabolic engineering. Trends in Microbiology 19, 323–329 (2011).

18. Kim, S.K. et al. A Genetically Encoded Biosensor for Monitoring Isoprene Production in Engineered Escherichia coli. ACS synthetic biology 7, 2379–2390 (2018).

19. Baojun, W., Mauricio, B. & Martin, B. Amplification of small molecule-inducible gene expression via tuning of intracellular receptor densities. Nucleic Acids Research 43, 1955–1964 (2015).

20. Srivatsan, R., Rogers, J.K., Taylor, N.D. & Church, G.M. Evolution-guided optimization of biosynthetic pathways. Proceedings of the National Academy of Sciences of the United States of America 111, 17803 (2014).

21. Ceroni, F. et al. Burden-driven feedback control of gene expression. Nature methods 15, 387 (2018).

22. Faure, G., Ogurtsov, A.Y., Shabalina, S.A. & Koonin, E.V. Role of mRNA structure in the control of protein folding. Nucleic acids research 44, 10898–10911 (2016).

23. Mutalik, V.K. et al. Precise and reliable gene expression via standard transcription and translation initiation elements. Nature Methods 10, 354 (2013).

24. Sauer, C. et al. Exploring the non-conserved sequence space of synthetic expression modules in Bacillus subtilis. ACS Synthetic Biology, acssynbio.8b00110- (2018).

25. Wang, J.D., Herman, C., Tipton, K.A., Gross, C.A. & Weissman, J.S. Directed evolution of substrate-optimized GroEL/S chaperonins. Cell 111, 1027–1039 (2002).

26. Crooks, G.E., Hon, G., Chandonia, J.-M. & Brenner, S.E. WebLogo: a sequence logo generator. Genome research 14, 1188–1190 (2004).

27. Nielsen, A.A., Segall-Shapiro, T.H. & Voigt, C.A. Advances in genetic circuit design: novel biochemistries, deep part mining, and precision gene expression. Current Opinion in Chemical Biology 17, 878–892 (2013).

28. Nielsen, A.A. et al. Genetic circuit design automation. Science 352, aac7341–aac7341 (2016).

29. Tae Seok, M., Chunbo, L., Alvin, T., Stanton, B.C. & Voigt, C.A. Genetic programs constructed from layered logic gates in single cells. Nature 491, 249–253 (2012).

30. Guo, J., Wang, T., Guan, C., Liu, B. & Xing, X.-H. Improved sgRNA design in bacteria via genome-wide activity profiling. Nucleic Acids Research 46, 7052–7069 (2018).

31. Zhou, S., Ding, R., Chen, J., Du, G. & Zhou, J. Obtaining a Panel of Cascade Promoter-5’-UTR Complexes in Escherichia coli. Acs Synthetic Biology 6, 1065–1075 (2017).

32. Sundaram, L. et al. Predicting the clinical impact of human mutation with deep neural networks. Nature Genetics 50 (2018).

33. Zhou, J. & Troyanskaya, O.G. Predicting effects of noncoding variants with deep learning– based sequence model. Nature methods 12, 931 (2015).

34. Gibson, D. et al. Enzymatic assembly of DNA molecules up to several hundred kilobases. Nature Methods 6, 343 (2009).

35. Jean-Denis, P., Stéphanie, C., Timothy, T., Terwilliger, T.C. & Waldo, G.S. Engineering and characterization of a superfolder green fluorescent protein. Nature Biotechnology 24, 79–88 (2006).

36. Jiang, Y. et al. Multigene editing in the Escherichia coli genome via the CRISPR-Cas9 system. Appl. Environ. Microbiol. 81, 2506–2514 (2015).

37. Ramanan, R.N. & Ariff, L.A.B. The Performance Of A Glass Bead Shaking Technique For The Disruption Of Escherichiacoli Cells. Biotechnology & Bioprocess Engineering 13, 613–623 (2008).

38. Liu, P., Wang, W., Zhao, J. & Wei, D. Screening novel β-galactosidases from a sequence-based metagenome and characterization of an alkaline β-galactosidase for the enzymatic synthesis of galactooligosaccharides. Protein expression and purification 155, 104–111 (2019).

39. Schaefer, J., Jovanovic, G., Kotta-Loizou, I. & Buck, M. Single-step method for β-galactosidase assays in Escherichia coli using a 96-well microplate reader. Analytical Biochemistry 503, 56–57 (2016).

