## Supplementary Materials for "Fine-tuning biosensor dynamic range based on rational design of cross-ribosome-binding sites in bacteria"

### Supplementary Figures

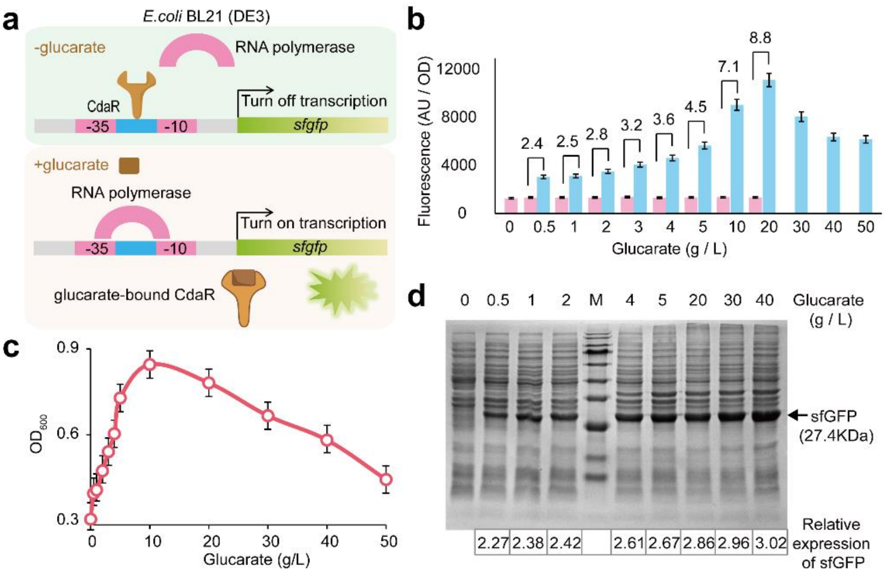


**Supplementary Fig. 1 Fluorescence intensity of glucarate biosensor in various glucarate concentration in vitro.** (**a**) The work principle of glucarate biosensor. In the absence of glucarate, CdaR binds to the CdaR-recognition site in a promoter, prevents RNA polymerase from binding to the promoter and represses *sfgfp* transcription. In the presence of glucarate, glucarate confronts the DNA-binding activity of CdaR, and RNA polymerase is able to bind to the promoter to turn on *sfgfp* transcription. (**b**) Fluorescence intensity and dynamic range of glucarate biosensor in diverse glucarate concentration. (**c**) Absorbance of glucarate biosensor-containing cells at 600 nm in various glucarate concentration. Cells were cultured in 96-well plates. (**d**) SDS-PAGE analysis of glucarate biosensor in diverse glucarate concentration. Relative expression of sfGFP was obtained by calculating the band density using ImageJ software.

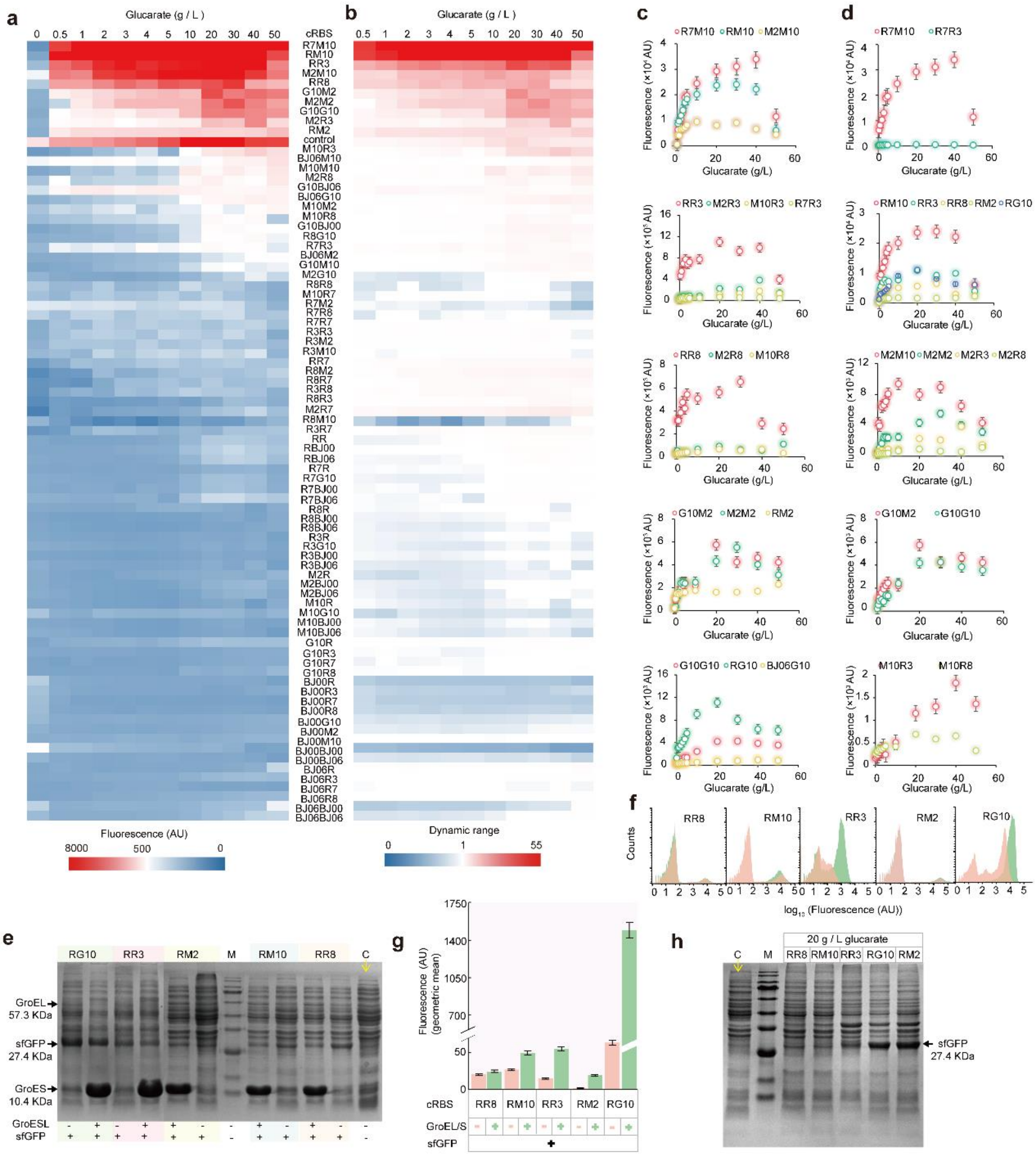


**Supplementary Fig. 2 Effects of cRBSs on biosensor dynamic range.** (**a**) The fluorescence intensity and (**b**) dynamic range of 81 cRBSs with various glucarate concentrations. (**c**) The effect of RBSn changes on fluorescence intensity under the same RBSm. (**d**) The effect of RBSm changes on fluorescence intensity under the same RBSn. (**e**) SDS-PAGE analysis of sfGFP, GroEL, GroES. For five cRBSs, GroEL/S overexpression had little effect on the level of GFP expression. Equivalent cell extracts of different cRBSs were analyzed by SDS-PAGE and visualized by Coomassie blue staining. The locations of GroEL, GroES and sfGFP are indicated. GroEL/S were expressed upon the addition of 1 mM IPTG. sfGFP was expression upon the addition of 20 g/L glucarate for each cRBSs. M: protein marker. C: control (BL21 (DE3)). (**f**) Expression strength analysis of sfGFP by FACS. The fluorescence intensity of sfGFP controlled by cRBSs were analyzed upon the addition of 20 g/L glucarate. (**g**) Analysis the fluorescence expression by flow cytometry. The geometric means corresponding to each cRBSs (light red and green columns) were calculated with and without GroEL/S (controlled by IPTG). (**h**) SDS-PAGE analysis of five cRBSs biosensors with 20 g / L glucarate. M: protein marker. C: control (BL21 (DE3)).

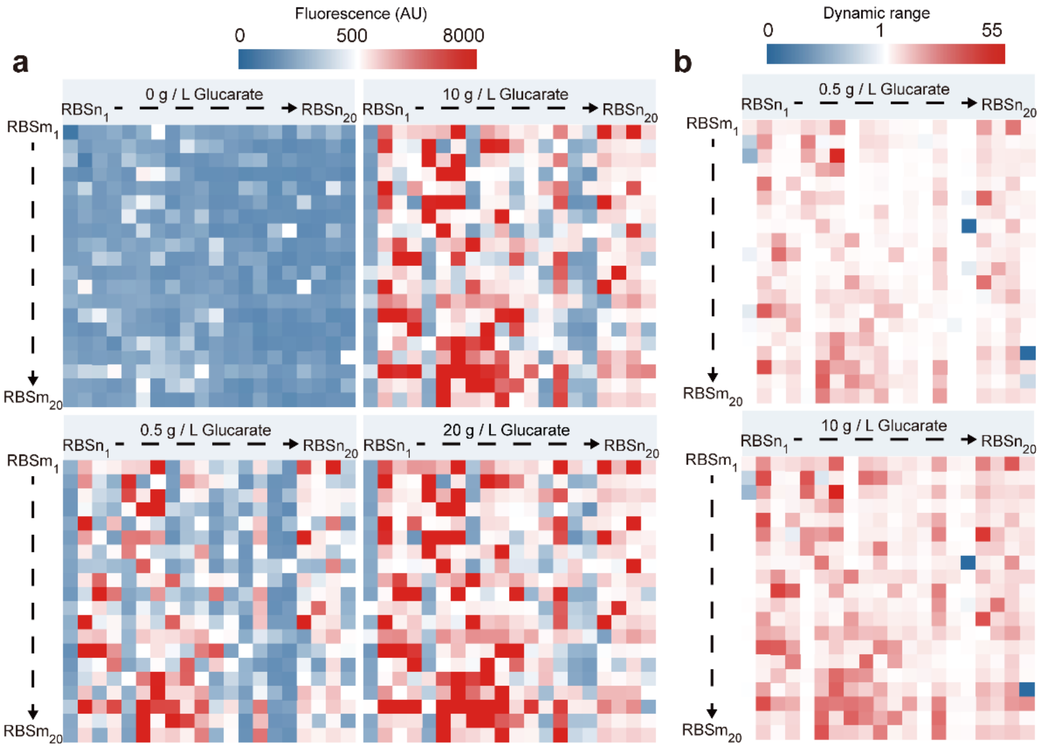


**Supplementary Fig. 3 Effects of semi-rational designed 400 cRBSs on biosensor fluorescence intensity and dynamic range.** (**a**) Comparisons of the fluorescence intensities of 400 cRBSs upon the addition of 0 g/L, 0.5 g/L, 10 g/L and 20 g/L glucarate. (**b**) The biosensor dynamic range of 400 cRBSs upon the addition of 0.5 g/L and 10 g/L glucarate. $RBSn_1$ – $RBSn_{20}$: RBS of *cdaR*; $RBSm_1$ – $RBSm_{20}$: RBS of *sfgfp*;

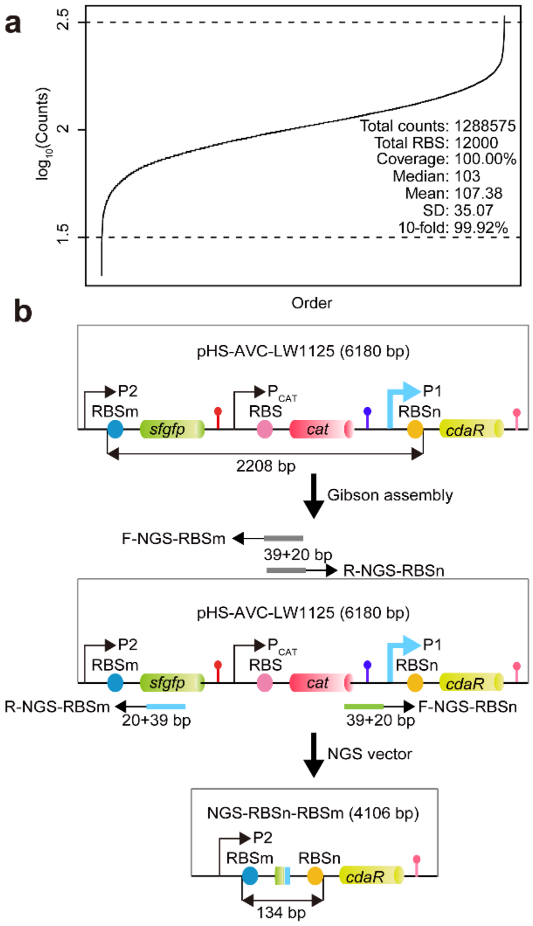


**Supplementary Fig. 4 NGS profiling of cRBS library in glucarate biosensor.** (**a**) Library profile at cRBS level presented by read count (log10) distribution of cRBSs detected in NGS. 10-fold variation is shown for distribution profile calculated by summarizing the event number (cRBS) with read count belonging to [1/3.33×median event read count, 3.33×median event read count], and normalized by the total event number. (**b**) Modifying the vector of five sub-libraries to further NGS analysis by gibson assembly.

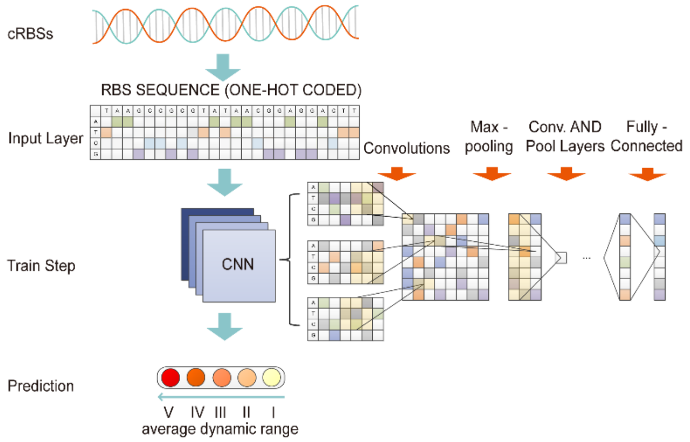


**Supplementary Fig. 5 Schematic overview of the CLM-RDR model.** 85% of the cRBSs datasets in each sub-library was used as input, the CNN was used as the training model, and ranks I–V divided by the average dynamic ranges of the glucarate biosensors were used as the model output. CNN model includes convolutions layers, max-pooling layers and fully-connected layers.

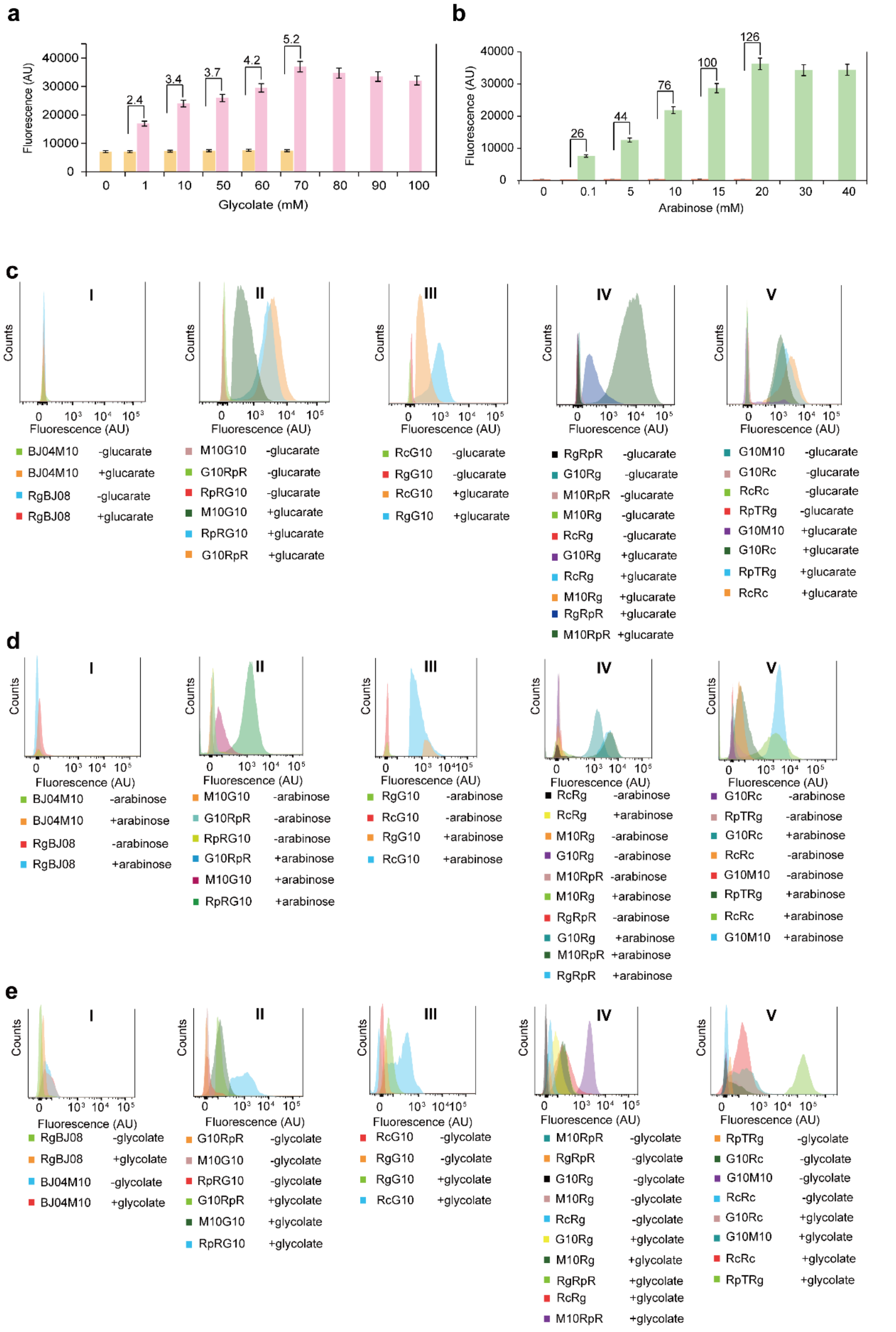


**Supplementary Fig. 6 Application of CLM-RDR model to other biosensors.** (**a**) Fluorescence intensity and dynamic range of glycolate biosensor in diverse glycolate concentration in vitro.

Glycolate biosensor had a dynamic range of up to 5.2 with 70 mM glycolate. (**b**) Fluorescence intensity and dynamic range of arabinose biosensor in various arabinose concentration in vitro. Arabinose biosensor had a dynamic range of up to 126 with 20 mM arabinose. (**c**) Comparison of Experimentally-observed fluorescence intensity of each cRBS with and without inducer in glucarate biosensor by Flow Cytometry. (**d**) Comparison of Experimentally-observed fluorescence intensity of each cRBS with and without inducer in arabinose biosensor by Flow Cytometry. (**e**) Comparison of Experimentally-observed fluorescence intensity of each cRBS with and without inducer in glycolate biosensor by Flow Cytometry. I-V represent prediction.

#### Supplementary Tables

**Supplementary Table 1 Performance for translation control element and inducer**

| | Sum of Squares | % | Mean Square | % | F | Sig. |
|---|---|---|---|---|---|---|
| cRBS | 12097752835.83 | 72.91 | 151221910.45 | 84.28 | 483.46 | 0.00E+00 |
| glucarate | 257666727.06 | 1.55 | 23424247.91 | 13.05 | 74.89 | 2.02E-121 |
| cRBS * glucarate | 3932946784.20 | 23.70 | 4469257.71 | 2.49 | 14.29 | 1.377E-294 |
| Error | 304034084.60 | 1.83 | 312792.27 | 0.17 | NA | NA |

ANOVA table for cRBSs elements and their interactions with inducer glucarate is shown at the table. Sum of squares represent the actual explanation of variation in the output measurement (fluorescence). The mean squares represent the average contribution of each of the factors/interactions taking into account their degrees of freedom (df).

**Supplementary Table 2 RBS strength analysis**

| | Translation Initiation Rate (au) targeting gene *cdaR* | ΔG_total (kcal mol-1) (*cdaR*) | Translation Initiation Rate (au) targeting gene *sfgfp* | ΔG_total (kcal mol-1) (*sfgfp*) | sequence（5' to 3'） |
|---|---|---|---|---|---|
| RBS | 137.46 | 4.88 | 36.57 | 9.39 | TAAGCCGTGCATAACGGAGGACTT |
| RBS3 | 4.78 | 12.34 | 127.10 | 5.05 | TTCCATTAAGAGGTAATTAAG |
| RBS7 | 81.59 | 6.03 | 155.29 | 4.6 | ATCCCATTCTTCTAGGAGTCGGC |
| RBS8 | 19.55 | 9.21 | 17.16 | 9.5 | GCAAAGAGGAGTTTAAACTTC |
| G10RBS | 2349.96 | -1.43 | 981.48 | 0.51 | TTTAACTTTAAGAAGGAGATATACAT |
| MCD2 | 7312.13 | -3.95 | 17428.75 | -5.88 | TCTTAATCATGCTAAGGAGGTTTTC |
| MCD10 | 310.43 | 3.07 | 437.03 | 2.31 | TCTTAATCATGCGGAGGATCGTTTC |
| BBa_J61100 | 313.53 | 3.04 | 194.58 | 4.10 | TCTAGAGAAAGAGGGGACAAACTAG |
| BBa_J61106 | 644.17 | 1.44 | 260.70 | 3.45 | TCTAGAGAAAGATAGGAGACACTAG |
| RBSn$_{81}$ | 190.43 | 4.15 | | | TAACCATGCATAAGGGAGCAGACTT |
| RBSm$_{56}$ | | | 995.86 | 0.48 | TCTTAATCATGGGAAGGAAGGTTTC |
| RBSm$_{97}$ | | | 1652.26 | -0.65 | TCTTAATCATGAGGAGGCTGGTTTC |
| RBSm$_{117}$ | | | 107.81 | 5.42 | TCTTAATCATGTGAAGAATGGTTTC |

**Supplementary Table 3 RBS sequence of *cdaR* and *sfgfp***

| RBS | Sequence |
|---|---|
| cdaR-RBSn$_1$ | TAACCATGCATAACGGAGTCGACTT |
| cdaR-RBSn$_2$ | TAACCATGCATATAGGAGGAGACTT |
| cdaR-RBSn$_3$ | TAACCATGCATAAAGGAGTCGACTT |

| | |
|---|---|
| cdaR-RBSn$_4$ | TAACCATGCATAGAGGAGCTGACTT |
| cdaR-RBSn$_5$ | TAACCATGCATAAAGGAGAAGACTT |
| cdaR-RBSn$_6$ | TAACCATGCATAAAGGAGGAGACTT |
| cdaR-RBSn$_7$ | TAACCATGCATAATGGAGGAGACTT |
| cdaR-RBSn$_8$ | TAACCATGCATAAGGGAGAGGACTT |
| cdaR-RBSn$_9$ | TAACCATGCATAAGGGAGGAGACTT |
| cdaR-RBSn$_{10}$ | TAACCATGCATATAGGAGAGGACTT |
| cdaR-RBSn$_{11}$ | TAACCATGCATATTGGAGAAGACTT |
| cdaR-RBSn$_{12}$ | TAACCATGCATATGGGAGAGGACTT |
| cdaR-RBSn$_{13}$ | TAACCATGCATATGGGAGTGGACTT |
| cdaR-RBSn$_{14}$ | TAACCATGCATACTGGAGGAGACTT |
| cdaR-RBSn$_{15}$ | TAACCATGCATACGGGAGATGACTT |
| cdaR-RBSn$_{16}$ | TAACCATGCATACGGGAGTGGACTT |
| cdaR-RBSn$_{17}$ | TAACCATGCATAGAGGAGAGGACTT |
| cdaR-RBSn$_{18}$ | TAACCATGCATAGAGGAGGTGACTT |
| cdaR-RBSn$_{19}$ | TAACCATGCATAGTGGAGGAGACTT |
| cdaR-RBSn$_{20}$ | TAACCATGCATAGGGGAGAGGACTT |
| cdaR-RBSn$_{21}$ | TAACCATGCATAAAGGAGATGACTT |
| cdaR-RBSn$_{22}$ | TAACCATGCATAAAGGAGACGACTT |
| cdaR-RBSn$_{23}$ | TAACCATGCATAAAGGAGAGGACTT |
| cdaR-RBSn$_{24}$ | TAACCATGCATAAAGGAGTAGACTT |
| cdaR-RBSn$_{25}$ | TAACCATGCATAAAGGAGTTGACTT |
| cdaR-RBSn$_{26}$ | TAACCATGCATAATGGAGAAGACTT |
| cdaR-RBSn$_{27}$ | TAACCATGCATAATGGAGATGACTT |
| cdaR-RBSn$_{28}$ | TAACCATGCATAATGGAGACGACTT |
| cdaR-RBSn$_{29}$ | TAACCATGCATATAGGAGAAGACTT |
| cdaR-RBSn$_{30}$ | TAACCATGCATATAGGAGATGACTT |
| cdaR-RBSn$_{31}$ | TAACCATGCATATAGGAGACGACTT |

| | |
|---|---|
| cdaR-RBSn$_{32}$ | TAACCATGCATAGAGGAGAAGACTT |
| cdaR-RBSn$_{33}$ | TAACCATGCATAGAGGAGATGACTT |
| cdaR-RBSn$_{34}$ | TAACCATGCATAGAGGAGACGACTT |
| cdaR-RBSn$_{35}$ | TAACCATGCATAGAGGAGGAGACTT |
| cdaR-RBSn$_{36}$ | TAACCATGCATAGTGGAGAAGACTT |
| cdaR-RBSn$_{37}$ | TAACCATGCATAGTGGAGATGACTT |
| cdaR-RBSn$_{38}$ | TAACCATGCATAGTGGAGACGACTT |
| cdaR-RBSn$_{39}$ | TAACCATGCATAGTGGAGAGGACTT |
| cdaR-RBSn$_{40}$ | TAACCATGCATAGGGGAGGAGACTT |
| cdaR-RBSn$_{41}$ | TAACCATGCATAAAGGAGTGGACTT |
| cdaR-RBSn$_{42}$ | TAACCATGCATAAAGGAGCAGACTT |
| cdaR-RBSn$_{43}$ | TAACCATGCATATGGGAGAAGACTT |
| cdaR-RBSn$_{44}$ | TAACCATGCATATGGGAGGAGACTT |
| cdaR-RBSn$_{45}$ | TAACCATGCATACAGGAGAGGACTT |
| cdaR-RBSn$_{46}$ | TAACCATGCATAAAGGAGGAGACTT |
| cdaR-RBSn$_{47}$ | TAACCATGCATAAAGGAGGCGACTT |
| cdaR-RBSn$_{48}$ | TAACCATGCATAATGGAGAGGACTT |
| cdaR-RBSn$_{49}$ | TAACCATGCATAATGGAGTAGACTT |
| cdaR-RBSn$_{50}$ | TAACCATGCATATGGGAGGTGACTT |
| cdaR-RBSn$_{51}$ | TAACCATGCATAATGGAGTCGACTT |
| cdaR-RBSn$_{52}$ | TAACCATGCATAATGGAGTGGACTT |
| cdaR-RBSn$_{53}$ | TAACCATGCATAATGGAGCAGACTT |
| cdaR-RBSn$_{54}$ | TAACCATGCATAATGGAGCTGACTT |
| cdaR-RBSn$_{55}$ | TAACCATGCATATGGGAGGCGACTT |
| cdaR-RBSn$_{56}$ | TAACCATGCATAATGGAGCGGACTT |
| cdaR-RBSn$_{57}$ | TAACCATGCATAATGGAGGTGACTT |
| cdaR-RBSn$_{58}$ | TAACCATGCATAATGGAGGCGACTT |
| cdaR-RBSn$_{59}$ | TAACCATGCATAACGGAGAAGACTT |

| | |
|---|---|
| cdaR-RBSn$_{60}$ | TAACCATGCATATGGGAGATGACTT |
| cdaR-RBSn$_{61}$ | TAACCATGCATATGGGAGACGACTT |
| cdaR-RBSn$_{62}$ | TAACCATGCATAACGGAGAGGACTT |
| cdaR-RBSn$_{63}$ | TAACCATGCATAGAGGAGGCGACTT |
| cdaR-RBSn$_{64}$ | TAACCATGCATAGGGGAGATGACTT |
| cdaR-RBSn$_{65}$ | TAACCATGCATAGGGGAGACGACTT |
| cdaR-RBSn$_{66}$ | TAACCATGCATACAGGAGGAGACTT |
| cdaR-RBSn$_{67}$ | TAACCATGCATAGGGGAGAAGACTT |
| cdaR-RBSn$_{68}$ | TAACCATGCATAAGGGAGGGGACTT |
| cdaR-RBSn$_{69}$ | TAACCATGCATATAGGAGGGGACTT |
| cdaR-RBSn$_{70}$ | TAACCATGCATAACGGAGGAGACTT |
| cdaR-RBSn$_{71}$ | TAACCATGCATAACGGAGGTGACTT |
| cdaR-RBSn$_{72}$ | TAACCATGCATAACGGAGGCGACTT |
| cdaR-RBSn$_{73}$ | TAACCATGCATAACGGAGGGGACTT |
| cdaR-RBSn$_{74}$ | TAACCATGCATAAGGGAGAAGACTT |
| cdaR-RBSn$_{75}$ | TAACCATGCATAAGGGAGATGACTT |
| cdaR-RBSn$_{76}$ | TAACCATGCATAAGGGAGACGACTT |
| cdaR-RBSn$_{77}$ | TAACCATGCATAAGGGAGTAGACTT |
| cdaR-RBSn$_{78}$ | TAACCATGCATAGAGGAGTAGACTT |
| cdaR-RBSn$_{79}$ | TAACCATGCATAAGGGAGTCGACTT |
| cdaR-RBSn$_{80}$ | TAACCATGCATAAGGGAGTGGACTT |
| cdaR-RBSn$_{81}$ | TAACCATGCATAAGGGAGCAGACTT |
| cdaR-RBSn$_{82}$ | TAACCATGCATAGAGGAGTTGACTT |
| cdaR-RBSn$_{83}$ | TAACCATGCATAGAGGAGTCGACTT |
| cdaR-RBSn$_{84}$ | TAACCATGCATAGAGGAGTGGACTT |
| cdaR-RBSn$_{85}$ | TAACCATGCATAAGGGAGGTGACTT |
| cdaR-RBSn$_{86}$ | TAACCATGCATAAGGGAGGCGACTT |
| cdaR-RBSn$_{87}$ | TAACCATGCATATAGGAGTAGACTT |

| | |
|---|---|
| cdaR-RBS$n_{88}$ | TAACCATGCATAGCGGAGAGGACTT |
| cdaR-RBS$n_{89}$ | TAACCATGCATAATGGAGGGGACTT |
| cdaR-RBS$n_{90}$ | TAACCATGCATATAGGAGTGGACTT |
| cdaR-RBS$n_{91}$ | TAACCATGCATACAGGAGAAGACTT |
| cdaR-RBS$n_{92}$ | TAACCATGCATACAGGAGATGACTT |
| cdaR-RBS$n_{93}$ | TAACCATGCATACAGGAGACGACTT |
| cdaR-RBS$n_{94}$ | TAACCATGCATATAGGAGCGGACTT |
| cdaR-RBS$n_{95}$ | TAACCATGCATATAGGAGGTGACTT |
| cdaR-RBS$n_{96}$ | TAACCATGCATATAGGAGGCGACTT |
| cdaR-RBS$n_{97}$ | TAACCATGCATATAGGAGGGGACTT |
| cdaR-RBS$n_{98}$ | TAACCATGCATATTGGAGATGACTT |
| cdaR-RBS$n_{99}$ | TAACCATGCATATTGGAGACGACTT |
| cdaR-RBS$n_{100}$ | TAACCATGCATATTGGAGAGGACTT |
| sfgfp-RBS$m_1$ | TCTTAATCATGCGGAGGAGGGTTTC |
| sfgfp-RBS$m_2$ | TCTTAATCATGAAGAGGATGGTTTC |
| sfgfp-RBS$m_3$ | TCTTAATCATGTCGAGCGAGGTTTC |
| sfgfp-RBS$m_4$ | TCTTAATCATGCTAAGAAGGGTTTC |
| sfgfp-RBS$m_5$ | TCTTAATCATGAGAAGAGTGGTTTC |
| sfgfp-RBS$m_6$ | TCTTAATCATGCAGAGGACGGTTTC |
| sfgfp-RBS$m_7$ | TCTTAATCATGTTGAGAGGGGTTTC |
| sfgfp-RBS$m_8$ | TCTTAATCATGTGAAGGCAGGTTTC |
| sfgfp-RBS$m_9$ | TCTTAATCATGCAAAGAGAGGTTTC |
| sfgfp-RBS$m_{10}$ | TCTTAATCATGGAGAGGTGGGTTTC |
| sfgfp-RBS$m_{11}$ | TCTTAATCATGGGAAGGATGGTTTC |
| sfgfp-RBS$m_{12}$ | TCTTAATCATGTAGAGGAAGGTTTC |
| sfgfp-RBS$m_{13}$ | TCTTAATCATGAGTAGGAGGGTTTC |
| sfgfp-RBS$m_{14}$ | TCTTAATCATGATGAGTGGGGTTTC |
| sfgfp-RBS$m_{15}$ | TCTTAATCATGCGAAGAATGGTTTC |

| | |
|---|---|
| sfgfp-RBS$m_{16}$ | TCTTAATCATGCTGAGAGCGGTTTC |
| sfgfp-RBS$m_{17}$ | TCTTAATCATGCTAAGAGAGGTTTC |
| sfgfp-RBS$m_{18}$ | TCTTAATCATGTGGAGGTAGGTTTC |
| sfgfp-RBS$m_{19}$ | TCTTAATCATGAAGAGTGAGGTTTC |
| sfgfp-RBS$m_{20}$ | TCTTAATCATGATAAGGCGGGTTTC |
| sfgfp-RBS$m_{21}$ | TCTTAATCATGAAGAGGAAGGTTTC |
| sfgfp-RBS$m_{22}$ | TCTTAATCATGAAGAGGACGGTTTC |
| sfgfp-RBS$m_{23}$ | TCTTAATCATGAAGAGGAGGGTTTC |
| sfgfp-RBS$m_{24}$ | TCTTAATCATGAGGAGGAAGGTTTC |
| sfgfp-RBS$m_{25}$ | TCTTAATCATGAGGAGGATGGTTTC |
| sfgfp-RBS$m_{26}$ | TCTTAATCATGAGGAGGACGGTTTC |
| sfgfp-RBS$m_{27}$ | TCTTAATCATGAGGAGGAGGGTTTC |
| sfgfp-RBS$m_{28}$ | TCTTAATCATGTGGAGGAAGGTTTC |
| sfgfp-RBS$m_{29}$ | TCTTAATCATGTGGAGGATGGTTTC |
| sfgfp-RBS$m_{30}$ | TCTTAATCATGTGGAGGACGGTTTC |
| sfgfp-RBS$m_{31}$ | TCTTAATCATGTGGAGGAGGGTTTC |
| sfgfp-RBS$m_{32}$ | TCTTAATCATGTAGAGGATGGTTTC |
| sfgfp-RBS$m_{33}$ | TCTTAATCATGTAGAGGACGGTTTC |
| sfgfp-RBS$m_{34}$ | TCTTAATCATGTAGAGGAGGGTTTC |
| sfgfp-RBS$m_{35}$ | TCTTAATCATGAGAAGGAAGGTTTC |
| sfgfp-RBS$m_{36}$ | TCTTAATCATGAGAAGGATGGTTTC |
| sfgfp-RBS$m_{37}$ | TCTTAATCATGAGAAGGACGGTTTC |
| sfgfp-RBS$m_{38}$ | TCTTAATCATGAGAAGGAGGGTTTC |
| sfgfp-RBS$m_{39}$ | TCTTAATCATGTGAAGGATGGTTTC |
| sfgfp-RBS$m_{40}$ | TCTTAATCATGTGAAGGAGGGTTTC |
| sfgfp-RBS$m_{41}$ | TCTTAATCATGAAGAGAAAGGTTTC |
| sfgfp-RBS$m_{42}$ | TCTTAATCATGAAGAGAATGGTTTC |
| sfgfp-RBS$m_{43}$ | TCTTAATCATGAAGAGAACGGTTTC |

| | |
|---|---|
| sfgfp-RBSm$_{44}$ | TCTTAATCATGAAGAGAAGGGTTTC |
| sfgfp-RBSm$_{45}$ | TCTTAATCATGAAGAGGCGGGTTTC |
| sfgfp-RBSm$_{46}$ | TCTTAATCATGAAGAGGGAGGTTTC |
| sfgfp-RBSm$_{47}$ | TCTTAATCATGAAGAGGGTGGTTTC |
| sfgfp-RBSm$_{48}$ | TCTTAATCATGGGGAGGAAGGTTTC |
| sfgfp-RBSm$_{49}$ | TCTTAATCATGGGGAGGATGGTTTC |
| sfgfp-RBSm$_{50}$ | TCTTAATCATGGGGAGGACGGTTTC |
| sfgfp-RBSm$_{51}$ | TCTTAATCATGGGGAGGAGGGTTTC |
| sfgfp-RBSm$_{52}$ | TCTTAATCATGGGGAGGTAGGTTTC |
| sfgfp-RBSm$_{53}$ | TCTTAATCATGAAGAGAGAGGTTTC |
| sfgfp-RBSm$_{54}$ | TCTTAATCATGAAGAGAGTGGTTTC |
| sfgfp-RBSm$_{55}$ | TCTTAATCATGACAAGGAGGGTTTC |
| sfgfp-RBSm$_{56}$ | TCTTAATCATGGGAAGGAAGGTTTC |
| sfgfp-RBSm$_{57}$ | TCTTAATCATGGGAAGGACGGTTTC |
| sfgfp-RBSm$_{58}$ | TCTTAATCATGGGAAGGAGGGTTTC |
| sfgfp-RBSm$_{59}$ | TCTTAATCATGGGAAGGTAGGTTTC |
| sfgfp-RBSm$_{60}$ | TCTTAATCATGTGAAGGAAGGTTTC |
| sfgfp-RBSm$_{61}$ | TCTTAATCATGAGAAGAATGGTTTC |
| sfgfp-RBSm$_{62}$ | TCTTAATCATGAGAAGAACGGTTTC |
| sfgfp-RBSm$_{63}$ | TCTTAATCATGAGAAGAAGGGTTTC |
| sfgfp-RBSm$_{64}$ | TCTTAATCATGAGAAGATAGGTTTC |
| sfgfp-RBSm$_{65}$ | TCTTAATCATGAGAAGATTGGTTTC |
| sfgfp-RBSm$_{66}$ | TCTTAATCATGAGAAGATCGGTTTC |
| sfgfp-RBSm$_{67}$ | TCTTAATCATGCGAAGAGAGGTTTC |
| sfgfp-RBSm$_{68}$ | TCTTAATCATGCGAAGAGTGGTTTC |
| sfgfp-RBSm$_{69}$ | TCTTAATCATGCGAAGAGCGGTTTC |
| sfgfp-RBSm$_{70}$ | TCTTAATCATGCGAAGAGGGGTTTC |
| sfgfp-RBSm$_{71}$ | TCTTAATCATGAGAAGACGGGTTTC |

| | |
|---|---|
| sfgfp-RBSm$_{72}$ | TCTTAATCATGAGAAGAGAGGTTTC |
| sfgfp-RBSm$_{73}$ | TCTTAATCATGAGAAGAGCGGTTTC |
| sfgfp-RBSm$_{74}$ | TCTTAATCATGAGGAGTGCGGTTTC |
| sfgfp-RBSm$_{75}$ | TCTTAATCATGAGGAGTGGGGTTTC |
| sfgfp-RBSm$_{76}$ | TCTTAATCATGATAAGGATGGTTTC |
| sfgfp-RBSm$_{77}$ | TCTTAATCATGATAAGGACGGTTTC |
| sfgfp-RBSm$_{78}$ | TCTTAATCATGATAAGGAGGGTTTC |
| sfgfp-RBSm$_{79}$ | TCTTAATCATGAGGAGCAGGGTTTC |
| sfgfp-RBSm$_{80}$ | TCTTAATCATGGGAAGACGGGTTTC |
| sfgfp-RBSm$_{81}$ | TCTTAATCATGGGAAGAGAGGTTTC |
| sfgfp-RBSm$_{82}$ | TCTTAATCATGGGAAGAGTGGTTTC |
| sfgfp-RBSm$_{83}$ | TCTTAATCATGGGAAGAGCGGTTTC |
| sfgfp-RBSm$_{84}$ | TCTTAATCATGGGAAGAGGGGTTTC |
| sfgfp-RBSm$_{85}$ | TCTTAATCATGTGAAGAGAGGTTTC |
| sfgfp-RBSm$_{86}$ | TCTTAATCATGTGAAGAGTGGTTTC |
| sfgfp-RBSm$_{87}$ | TCTTAATCATGTGAAGAGCGGTTTC |
| sfgfp-RBSm$_{88}$ | TCTTAATCATGTGAAGAGGGGTTTC |
| sfgfp-RBSm$_{89}$ | TCTTAATCATGAGAAGGGAGGTTTC |
| sfgfp-RBSm$_{90}$ | TCTTAATCATGAGAAGGGTGGTTTC |
| sfgfp-RBSm$_{91}$ | TCTTAATCATGAGAAGGGCGGTTTC |
| sfgfp-RBSm$_{92}$ | TCTTAATCATGAGGAGGTAGGTTTC |
| sfgfp-RBSm$_{93}$ | TCTTAATCATGAGGAGGTTGGTTTC |
| sfgfp-RBSm$_{94}$ | TCTTAATCATGAGGAGGTCGGTTTC |
| sfgfp-RBSm$_{95}$ | TCTTAATCATGAGGAGGTGGGTTTC |
| sfgfp-RBSm$_{96}$ | TCTTAATCATGAGGAGGCAGGTTTC |
| sfgfp-RBSm$_{97}$ | TCTTAATCATGAGGAGGCTGGTTTC |
| sfgfp-RBSm$_{98}$ | TCTTAATCATGAGGAGGCCGGTTTC |
| sfgfp-RBSm$_{99}$ | TCTTAATCATGAGGAGGCGGGTTTC |

| | |
|---|---|
| sfgfp-RBSm$_{100}$ | TCTTAATCATGAGGAGGGAGGTTTC |
| sfgfp-RBSm$_{101}$ | TCTTAATCATGAGGAGGGTGGTTTC |
| sfgfp-RBSm$_{102}$ | TCTTAATCATGAGGAGGGCGGTTTC |
| sfgfp-RBSm$_{103}$ | TCTTAATCATGAAGAGAGCGGTTTC |
| sfgfp-RBSm$_{104}$ | TCTTAATCATGAAGAGAGGGGTTTC |
| sfgfp-RBSm$_{105}$ | TCTTAATCATGAAGAGTAGGGTTTC |
| sfgfp-RBSm$_{106}$ | TCTTAATCATGTGGAGGGAGGTTTC |
| sfgfp-RBSm$_{107}$ | TCTTAATCATGTGGAGGGTGGTTTC |
| sfgfp-RBSm$_{108}$ | TCTTAATCATGTGGAGGGCGGTTTC |
| sfgfp-RBSm$_{109}$ | TCTTAATCATGTGGAGGGGGGTTTC |
| sfgfp-RBSm$_{110}$ | TCTTAATCATGCGGAGAATGGTTTC |
| sfgfp-RBSm$_{111}$ | TCTTAATCATGCGGAGAACGGTTTC |
| sfgfp-RBSm$_{112}$ | TCTTAATCATGCGGAGAAGGGTTTC |
| sfgfp-RBSm$_{113}$ | TCTTAATCATGCGGAGATAGGTTTC |
| sfgfp-RBSm$_{114}$ | TCTTAATCATGTAGAGAATGGTTTC |
| sfgfp-RBSm$_{115}$ | TCTTAATCATGTAGAGAAGGGTTTC |
| sfgfp-RBSm$_{116}$ | TCTTAATCATGTGAAGAAAGGTTTC |
| sfgfp-RBSm$_{117}$ | TCTTAATCATGTGAAGAATGGTTTC |
| sfgfp-RBSm$_{118}$ | TCTTAATCATGTGAAGAACGGTTTC |
| sfgfp-RBSm$_{119}$ | TCTTAATCATGTGAAGAAGGGTTTC |
| sfgfp-RBSm$_{120}$ | TCTTAATCATGGGAAGAAGGGTTTC |

**Supplementary Table 4 Strains and plasmids used in this study**

| **Name** | **Relevant genotype** | **Reference** |
|---|---|---|
| JM109 | *recA*1, *endA*1, *gyrA*96, *thi*,*hsdR*17, *supE*44 , *relA*1,Δ *(lac*$^{-}$ | Prof. Zhou |

| | | |
|---|---|---|
| | *proAB*)/F'[*traD*36, *proAB*$^+$, *lacI*$^q$, *lacZ*ΔM 1] | |
| BL21(DE3) | F$^-$ *ompT hsdS*$_B$（$r_B^-$ $m_B^-$），*gal*，*dcm*（DE3） | Prof. Zhou |
| BL21(DE3) Δ*lacZ* | F$^-$ *ompT hsdS*$_B$（$r_B^-$ $m_B^-$），*gal*，*dcm*（DE3） Δ*lacZ* | This study |
| pJKR-H-RBS-*cdaR*-g10RBS | sfGFP, CdaR, Amp$^R$ | Addgene (#62557) |
| pJKR-H-RBSs-*cdaR*-RBSs | RBSs = R, R3, R7, R8, G10, M2, M10, BJ00, BJ06, Amp$^R$ | This study |
| pHS-AVC-LW1125 | sfGFP, CdaR, CAT, *cdaR*-SD mix(100), *sfgfp*-SD mix(120), Amp$^R$ | This study |
| NGS-RBSn-RBSm | *cdaR*-SD mix(100), *sfgfp*-SD mix(120), Amp$^R$ | This study |
| NGS-RBSn-RBSm-I | sub-libraries of plasmid of extremely low fluorescence intensity in sample (with glucarate), Amp$^R$ | This study |
| NGS-RBSn-RBSm-II | sub-libraries of plasmid of low fluorescence intensity in sample (with glucarate), Amp$^R$ | This study |
| NGS-RBSn-RBSm-III | sub-libraries of plasmid of medium fluorescence intensity in sample (with glucarate), Amp$^R$ | This study |
| NGS-RBSn-RBSm-IV | sub-libraries of plasmid of high fluorescence intensity in sample (with glucarate), Amp$^R$ | |
| NGS-RBSn-RBSm-V | sub-libraries of plasmid of extremely high fluorescence intensity in sample (with glucarate), Amp$^R$ | This study |

| | | |
|---|---|---|
| pCas | paraB-gam-bet-exo, bla ($kan^R$), kanR-repA101 (ts),λ-Red, the sgRNA with a $lacI^q$-Ptrc promoter guiding the pMB1 replication of pTarget | Addgene |
| pTarget | the sgRNA sequence, a targeting N20 sequence, the pMB1 replicon, $Spe^R$ | Addgene |
| pUC-*glcC*-ffs | sfGFP, GlcC, ffs, $Amp^R$ | This study |
| pUC-*araC* | sfGFP, AraC, $Amp^R$ | This study |
| pJKR-H-M10-*cdaR*-BJ04 | MCD10 (M10), BBa_J61104 (BJ04), $Amp^R$ | This study |
| pJKR-H-$R_c$-*cdaR*-$R_c$ | $RBS_{cdaR}$ ( $R_c$), $Amp^R$ | This study |
| pJKR-H- $R_g$ -*cdaR*- $R_c$ | $RBS_{glcC}$ ( $R_g$), $Amp^R$ | This study |
| pJKR-H- G10-*cdaR*- $R_c$ | G10RBS (G10), $Amp^R$ | This study |
| pJKR-H-BJ08-*cdaR*- $R_g$ | BBa_J61108 (BJ08), $Amp^R$ | This study |
| pJKR-H-$R_{pR}$-*cdaR*- $R_g$ | $RBS_{pRSF}$ ($R_{pR}$), $Amp^R$ | This study |
| pJKR-H-G10-*cdaR*- $R_g$ | $Amp^R$ | This study |
| pJKR-H-G10-*cdaR*- $R_{pR}$ | $Amp^R$ | This study |
| pJKR-H- $R_g$ -*cdaR*- $R_{pT}$ | $RBS_{pTrc99a}$ ($R_{pT}$), $Amp^R$ | This study |
| pJKR-H- $R_g$ -*cdaR*-M10 | $Amp^R$ | This study |
| pJKR-H- $R_{pR}$ -*cdaR*- M10 | $Amp^R$ | This study |
| pJKR-H-G10-*cdaR*- M10 | $Amp^R$ | This study |
| pJKR-H- $R_c$ -*cdaR*-G10 | $Amp^R$ | This study |
| pJKR-H- $R_g$ -*cdaR*- G10 | $Amp^R$ | This study |
| pJKR-H- $R_{pR}$ -*cdaR*-G10 | $Amp^R$ | This study |
| pJKR-H-M10 -*cdaR*-G10 | $Amp^R$ | This study |
| pUC-M10-*glcC*-ffs-BJ04 | $Amp^R$ | This study |
| pUC -$R_c$- *glcC*-ffs -$R_c$ | $Amp^R$ | This study |
| pUC - $R_g$ - *glcC*-ffs - $R_c$ | $Amp^R$ | This study |

| | | |
|---|---|---|
| pUC - G10- *glcC*-ffs - $R_c$ | $Amp^R$ | This study |
| pUC -BJ08- *glcC*-ffs - $R_g$ | $Amp^R$ | This study |
| pUC -$R_{pR}$- *glcC*-ffs - $R_g$ | $Amp^R$ | This study |
| pUC -G10- *glcC*-ffs - $R_g$ | $Amp^R$ | This study |
| pUC -G10- *glcC*-ffs - $R_{pR}$ | $Amp^R$ | This study |
| pUC - $R_g$ - *glcC*-ffs - $R_{pT}$ | $Amp^R$ | This study |
| pUC - $R_g$ - *glcC*-ffs -M10 | $Amp^R$ | This study |
| pUC - $R_{pR}$ -*glcC*-ffs - M10 | $Amp^R$ | This study |
| pUC -G10-*glcC*-ffs - M10 | $Amp^R$ | This study |
| pUC - $R_c$ - *glcC*-ffs -G10 | $Amp^R$ | This study |
| pUC - $R_g$ - *glcC*-ffs - G10 | $Amp^R$ | This study |
| pUC - $R_{pR}$ - *glcC*-ffs -G10 | $Amp^R$ | This study |
| pUC -M10 -*glcC*-ffs -G10 | $Amp^R$ | This study |
| pUC-M10-*araC*-BJ04 | $Amp^R$ | This study |
| pUC -$R_c$- *araC* -$R_c$ | $Amp^R$ | This study |
| pUC - $R_g$ - *araC* - $R_c$ | $Amp^R$ | This study |
| pUC - G10- *araC* - $R_c$ | $Amp^R$ | This study |
| pUC -BJ08- *araC* - $R_g$ | $Amp^R$ | This study |
| pUC -$R_{pR}$- *araC* - $R_g$ | $Amp^R$ | This study |
| pUC -G10- *araC* - $R_g$ | $Amp^R$ | This study |
| pUC -G10- *araC* - $R_{pR}$ | $Amp^R$ | This study |
| pUC - $R_g$ - *araC* - $R_{pT}$ | $Amp^R$ | This study |
| pUC - $R_g$ - *araC* -M10 | $Amp^R$ | This study |
| pUC - $R_{pR}$ - *araC* - M10 | $Amp^R$ | This study |
| pUC -G10- *araC* - M10 | $Amp^R$ | This study |
| pUC - $R_c$ - *araC* -G10 | $Amp^R$ | This study |
| pUC - $R_g$ - *araC* - G10 | $Amp^R$ | This study |
| pUC - $R_{pR}$ - *araC* -G10 | $Amp^R$ | This study |

| | | |
|---|---|---|
| pUC -M10 - *araC* -G10 | Amp$^{R}$ | This study |
| pRSF-*groEL* -*groES* | Kan$^{R}$ | This study |
| pJKR-H-RBS$_{81}$-*cdaR*- RBS$_{56}$ | Amp$^{R}$ | This study |
| pJKR-H-RBS$_{81}$-*cdaR*- RBS$_{97}$ | Amp$^{R}$ | This study |
| pJKR-H-RBS$_{81}$-*cdaR*- RBS$_{117}$ | Amp$^{R}$ | This study |

**Supplementary Table 5 Primers used in this study**

| Primers | Sequence (from 5' to 3') |
|---|---|
| F-RBS-GFP | TAAGCCGTGCATAACGGAGGACTTATGCGTATAGGTGAAGAACTG |
| R-RBS-GFP | AAGTCCTCCGTTATGCACGGCTTATGTTGCACTCCTGAAAATTCG |
| F-G10-cdaR | TTTAACTTTAAGAAGGAGATATACATATGGCTGGCTGGCATCTTGATACCAAAATGGCGC |
| R-G10-cdaR | GCCAGCCATATGTATATCTCCTTCTTAAAGTTAAAGCATAAGGAAGTACGTAACGTACGG |
| F-BJ00-GFP | TCTAGAGAAAGAGGGGACAAACTAGATGCGTATAGGTGAAGAACT |
| R-BJ00-GFP | CTAGTTTGTCCCCTCTTTCTCTAGATGTTGCACTCCTGAAAATTC |
| F-BJ00-cdaR | TCTAGAGAAAGAGGGGACAAACTAGATGGCTGGCTGGCATCTTGA |
| R-BJ00-cdaR | CTAGTTTGTCCCCTCTTTCTCTAGAGCATAAGGAAGTACGTAACG |
| F-BJ06-GFP | TCTAGAGAAAGATAGGAGACACTAGATGCGTATAGGTGAAGAACT |
| R-BJ06-GFP | CTAGTGTCTCCTATCTTTCTCTAGATGTTGCACTCCTGAAAATTC |
| F-BJ06-cdaR | TCTAGAGAAAGATAGGAGACACTAGATGGCTGGCTGGCATCTTGA |
| R-BJ06-cdaR | CTAGTGTCTCCTATCTTTCTCTAGAGCATAAGGAAGTACGTAACG |
| F-MCD2-GFP | TCTTAATCATGCTAAGGAGGTTTTCATGCGTATAGGTGAAGAACTGTTCACCGGTG |
| R-MCD2-GFP | GAAAACCTCCTTAGCATGATTAAGATGTTGCACTCCTGAAAATTCGCGTTAGCCAC |

| | |
|---|---|
| F-MCD2-cdaR | TCTTAATCATGCTAAGGAGGTTTTCATGGCTGGCTGGCATCTTGATACCAAAATGG |
| R-MCD2-cdaR | GAAAACCTCCTTAGCATGATTAAGAGCATAAGGAAGTACGTAACGTACGGCATTGT |
| F-MCD10-GFP | TCTTAATCATGCGGAGGATCGTTTCATGCGTATAGGTGAAGAACTGTTCACCGGTG |
| R-MCD10-GFP | GAAACGATCCTCCGCATGATTAAGATGTTGCACTCCTGAAAATTCGCGTTAGCCAC |
| F-MCD10-cdaR | TCTTAATCATGCGGAGGATCGTTTCATGGCTGGCTGGCATCTTGATACCAAAATGG |
| R-MCD10-cdaR | GAAACGATCCTCCGCATGATTAAGAGCATAAGGAAGTACGTAACGTACGGCATTGT |
| F-RBS7-GFP | ATCCCATTCTTCTAGGAGTCGGCATGCGTATAGGTGAAGAACTGTTCACCGGTGT |
| R-RBS7-GFP | GCCGACTCCTAGAAGAATGGGATTGTTGCACTCCTGAAAATTCGCGTTAGCCACG |
| F-RBS7-cdaR | ATCCCATTCTTCTAGGAGTCGGCATGGCTGGCTGGCATCTTGATACCAAAATGGC |
| R-RBS7-cdaR | GCCGACTCCTAGAAGAATGGGATGCATAAGGAAGTACGTAACGTACGGCATTGTG |
| F-RBS8-GFP | GCAAAGAGGAGTTTAAACTTCATGCGTATAGGTGAAGAACTGTTCACCGGTGTTGTTC |
| R-RBS8-GFP | GAAGTTTAAACTCCTCTTTGCTGTTGCACTCCTGAAAATTCGCGTTAGCCACGCT |
| F-RBS8-cdaR | GCAAAGAGGAGTTTAAACTTCATGGCTGGCTGGCATCTTGATACCAAAATGGCGC |
| R-RBS8-cdaR | GAAGTTTAAACTCCTCTTTGCGCATAAGGAAGTACGTAACGTACGGCATTGTGC |

| | |
|---|---|
| F-RBS3-cdaR | TTCCATTAAGAGGTAATTAAGATGGCTGGCTGGCATCTTGATAC CAAAATGGCGC |
| R-RBS3-cdaR | CTTAATTACCTCTTAATGGAAGCATAAGGAAGTACGTAACGTAC GGCATTGTGC |
| F-lacZ | ATGACCATGATTACGGATTCACTGG |
| R-lacZ | TTATTTTTGACACCAGACCAACTGG |
| F-ZT | GGTCGCTACCATTACCAGTTGGTCTGGTGTCAAAAATAAATGCA TGCCAGTTCTAGCAT |
| R-G10ZT | CGTTGTAAAACGACGGCCAGTGAATCCGTAATCATGGTCATAT GTATATCTCCTTCTTA |
| R-R8ZT | TTGTAAAACGACGGCCAGTGAATCCGTAATCATGGTCATGAAG TTTAAACTCCTCTTTG |
| R-M10ZT | TTGTAAAACGACGGCCAGTGAATCCGTAATCATGGTCATGAAA CGATCCTCCGCATGAT |
| F-up | GGGCAGCAGCCATCACCATCATCACCACAGCCAGGATCCGGCA TCGTTCCCACTGCGAT |
| R-up | TACGCGAAATACGGGCAGACATGGCCTGCCCGGTTATTAAGCT GTTTCCTGTGTGAAAT |
| F-down | AAATTGTGAGCGGATAACAATTTCACACAGGAAACAGCTTAAT AACCGGGCAGGCCATG |
| R-down | ACTTTCTGTTCGACTTAAGCATTATGCGGCCGCAAGCTTCCAAC ACAGCCAAACATCCG |
| F-yzqc | GGCATCGTTCCCACTGCGAT |
| R-yzqc | CCAACACAGCCAAACATCCG |
| yz-F-SD | GCGGTGCTGAATCGAATC |
| yz-R-SD | ACACCGTGCAGATGACGT |
| FcdaR-RBS$n_1$ | TAACCATGCATAACGGAGTCGACTTATGGCTGGCTGGCATCTTG A |

| | |
|---|---|
| RcdaR-RBS$n_1$ | AAGTCGACTCCGTTATGCATGGTTAGCATAAGGAAGTACGTAACG |
| Fgfp-RBS$m_1$ | TCTTAATCATGCGGAGGAGGGTTTCATGCGTATAGGTGAAGAACT |
| Rgfp-RBS$m_1$ | GAAACCCTCCTCCGCATGATTAAGATGTTGCACTCCTGAAAATTC |
| FcdaR-RBS$n_2$ | TAACCATGCATATAGGAGGAGACTTATGGCTGGCTGGCATCTTGA |
| RcdaR-RBS$n_2$ | AAGTCTCCTCCTATATGCATGGTTAGCATAAGGAAGTACGTAACG |
| Fgfp-RBS$m_2$ | TCTTAATCATGAAGAGGATGGTTTCATGCGTATAGGTGAAGAACTGTTCACC |
| Rgfp-RBS$m_2$ | GAAACCATCCTCTTCATGATTAAGATGTTGCACTCCTGAAAATTCGCGTTAG |
| FcdaR-RBS$n_3$ | TAACCATGCATAAAGGAGTCGACTTATGGCTGGCTGGCATCTTGA |
| RcdaR-RBS$n_3$ | AAGTCGACTCCTTTATGCATGGTTAGCATAAGGAAGTACGTAACG |
| Fgfp-RBS$m_3$ | TCTTAATCATGTCGAGCGAGGTTTCATGCGTATAGGTGAAGAACT |
| Rgfp-RBS$m_3$ | GAAACCTCGCTCGACATGATTAAGATGTTGCACTCCTGAAAATTC |
| FcdaR-RBS$n_4$ | TAACCATGCATAGAGGAGCTGACTTATGGCTGGCTGGCATCTTGA |
| RcdaR-RBS$n_4$ | AAGTCAGCTCCTCTATGCATGGTTAGCATAAGGAAGTACGTAACG |
| Fgfp-RBS$m_4$ | TCTTAATCATGCTAAGAAGGGTTTCATGCGTATAGGTGAAGAACTGTTCACC |

| | |
|---|---|
| Rgfp-RBSm$_4$ | GAAACCCTTCTTAGCATGATTAAGATGTTGCACTCCTGAAAATTCGCGTTAG |
| FcdaR-RBSn$_5$ | TAACCATGCATAAAGGAGAAGACTTATGGCTGGCTGGCATCTT |
| RcdaR-RBSn$_5$ | AAGTCTTCTCCTTTATGCATGGTTAGCATAAGGAAGTACGTAACG |
| Fgfp-RBSm$_5$ | TCTTAATCATGAGAAGAGTGGTTTCATGCGTATAGGTGAAG |
| Rgfp-RBSm$_5$ | GAAACCACTCTTCTCATGATTAAGATGTTGCACTCCTGAA |
| FcdaR-RBSn$_6$ | TAACCATGCATAAAGGAGGAGACTTATGGCTGGCTGGCATCT |
| RcdaR-RBSn$_6$ | AAGTCTCCTCCTTTATGCATGGTTAGCATAAGGAAGTACGTA |
| Fgfp-RBSm$_6$ | TCTTAATCATGCAGAGGACGGTTTCATGCGTATAGGTGAAGA |
| Rgfp-RBSm$_6$ | GAAACCGTCCTCTGCATGATTAAGATGTTGCACTCCTGAAAAT |
| FcdaR-RBSn$_7$ | TAACCATGCATAATGGAGGAGACTTATGGCTGGCTGGCATCTT |
| RcdaR-RBSn$_7$ | AAGTCTCCTCCATTATGCATGGTTAGCATAAGGAAGTACGTAAC |
| Fgfp-RBSm$_7$ | TCTTAATCATGTTGAGAGGGGTTTCATGCGTATAGGTGAAGAAC |
| Rgfp-RBSm$_7$ | GAAACCCCTCTCAACATGATTAAGATGTTGCACTCCTGAAAATTC |
| FcdaR-RBSn$_8$ | TAACCATGCATAAGGGAGAGGACTTATGGCTGGCTGGCATCTT |
| RcdaR-RBSn$_8$ | AAGTCCTCTCCCTTATGCATGGTTAGCATAAGGAAGTACGTAA |
| Fgfp-RBSm$_8$ | TCTTAATCATGTGAAGGCAGGTTTCATGCGTATAGGTGAAGAAC |
| Rgfp-RBSm$_8$ | GAAACCTGCCTTCACATGATTAAGATGTTGCACTCCTGAAAATTC |
| FcdaR-RBSn$_9$ | TAACCATGCATAAGGGAGGAGACTTATGGCTGGCTGGCATCTT |
| RcdaR-RBSn$_9$ | AAGTCTCCTCCCTTATGCATGGTTAGCATAAGGAAGTACGTAA |
| Fgfp-RBSm$_9$ | TCTTAATCATGCAAAGAGAGGTTTCATGCGTATAGGTGAAG |
| Rgfp-RBSm$_9$ | GAAACCTCTCTTTGCATGATTAAGATGTTGCACTCCTGAA |
| FcdaR-RBSn$_{10}$ | TAACCATGCATATAGGAGAGGACTTATGGCTGGCTGGCAT |
| RcdaR-RBSn$_{10}$ | AAGTCCTCTCCTATATGCATGGTTAGCATAAGGAAGTACG |
| Fgfp-RBSm$_{10}$ | TCTTAATCATGGAGAGGTGGGTTTCATGCGTATAGGTGAA |

| | |
|---|---|
| Rgfp-RBSm$_{10}$ | GAAACCCACCTCTCCATGATTAAGATGTTGCACTCCTGAA |
| FcdaR-RBSn$_{11}$ | TAACCATGCATATTGGAGAAGACTTATGGCTGGCTGGCAT |
| RcdaR-RBSn$_{11}$ | AAGTCTTCTCCAATATGCATGGTTAGCATAAGGAAGTACG |
| Fgfp-RBSm$_{11}$ | TCTTAATCATGGGAAGGATGGTTTCATGCGTATAGGTGAA |
| Rgfp-RBSm$_{11}$ | GAAACCATCCTTCCCATGATTAAGATGTTGCACTCCTGAA |
| FcdaR-RBSn$_{12}$ | TAACCATGCATATGGGAGAGGACTTATGGCTGGCTGGCAT |
| RcdaR-RBSn$_{12}$ | AAGTCCTCTCCCATATGCATGGTTAGCATAAGGAAGTACG |
| Fgfp-RBSm$_{12}$ | TCTTAATCATGTAGAGGAAGGTTTCATGCGTATAGGTGAAG |
| Rgfp-RBSm$_{12}$ | GAAACCTTCCTCTACATGATTAAGATGTTGCACTCCTGAA |
| FcdaR-RBSn$_{13}$ | TAACCATGCATATGGGAGTGGACTTATGGCTGGCTGGCAT |
| RcdaR-RBSn$_{13}$ | AAGTCCACTCCCATATGCATGGTTAGCATAAGGAAGTACG |
| Fgfp-RBSm$_{13}$ | TCTTAATCATGAGTAGGAGGGTTTCATGCGTATAGGTGAA |
| Rgfp-RBSm$_{13}$ | GAAACCCTCCTACTCATGATTAAGATGTTGCACTCCTGAA |
| FcdaR-RBSn$_{14}$ | TAACCATGCATACTGGAGGAGACTTATGGCTGGCTGGCAT |
| RcdaR-RBSn$_{14}$ | AAGTCTCCTCCAGTATGCATGGTTAGCATAAGGAAGTACG |
| Fgfp-RBSm$_{14}$ | TCTTAATCATGATGAGTGGGGTTTCATGCGTATAGGTGAA |
| Rgfp-RBSm$_{14}$ | GAAACCCCACTCATCATGATTAAGATGTTGCACTCCTGAA |
| FcdaR-RBSn$_{15}$ | TAACCATGCATACGGGAGATGACTTATGGCTGGCTGGCAT |
| RcdaR-RBSn$_{15}$ | AAGTCATCTCCCGTATGCATGGTTAGCATAAGGAAGTACG |
| Fgfp-RBSm$_{15}$ | TCTTAATCATGCGAAGAATGGTTTCATGCGTATAGGTGAAG |
| Rgfp-RBSm$_{15}$ | GAAACCATTCTTCGCATGATTAAGATGTTGCACTCCTGAA |
| FcdaR-RBSn$_{16}$ | TAACCATGCATACGGGAGTGGACTTATGGCTGGCTGGCAT |
| RcdaR-RBSn$_{16}$ | AAGTCCACTCCCGTATGCATGGTTAGCATAAGGAAGTACG |
| Fgfp-RBSm$_{16}$ | TCTTAATCATGCTGAGAGCGGTTTCATGCGTATAGGTGAA |
| Rgfp-RBSm$_{16}$ | GAAACCGCTCTCAGCATGATTAAGATGTTGCACTCCTGAA |
| FcdaR-RBSn$_{17}$ | TAACCATGCATAGAGGAGAGGACTTATGGCTGGCTGGCAT |
| RcdaR-RBSn$_{17}$ | AAGTCCTCTCCTCTATGCATGGTTAGCATAAGGAAGTACG |
| Fgfp-RBSm$_{17}$ | TCTTAATCATGCTAAGAGAGGTTTCATGCGTATAGGTGAAG |

| | |
|---|---|
| Rgfp-RBSm$_{17}$ | GAAACCTCTCTTAGCATGATTAAGATGTTGCACTCCTGAA |
| FcdaR-RBSn$_{18}$ | TAACCATGCATAGAGGAGGTGACTTATGGCTGGCTGGCAT |
| RcdaR-RBSn$_{18}$ | AAGTCACCTCCTCTATGCATGGTTAGCATAAGGAAGTACG |
| Fgfp-RBSm$_{18}$ | TCTTAATCATGTGGAGGTAGGTTTCATGCGTATAGGTGAA |
| Rgfp-RBSm$_{18}$ | GAAACCTACCTCCACATGATTAAGATGTTGCACTCCTGAA |
| FcdaR-RBSn$_{19}$ | TAACCATGCATAGTGGAGGAGACTTATGGCTGGCTGGCAT |
| RcdaR-RBSn$_{19}$ | AAGTCTCCTCCACTATGCATGGTTAGCATAAGGAAGTACG |
| Fgfp-RBSm$_{19}$ | TCTTAATCATGAAGAGTGAGGTTTCATGCGTATAGGTGAAG |
| Rgfp-RBSm$_{19}$ | GAAACCTCACTCTTCATGATTAAGATGTTGCACTCCTGAA |
| FcdaR-RBSn$_{20}$ | TAACCATGCATAGGGGAGAGGACTTATGGCTGGCTGGCAT |
| RcdaR-RBSn$_{20}$ | AAGTCCTCTCCCCTATGCATGGTTAGCATAAGGAAGTACG |
| Fgfp-RBSm$_{20}$ | TCTTAATCATGATAAGGCGGGTTTCATGCGTATAGGTGAA |
| Rgfp-RBSm$_{20}$ | GAAACCCGCCTTATCATGATTAAGATGTTGCACTCCTGAA |
| F-NGS-RBSn | TTCACCGGTGTTGTTCCGATCTAGGCATTTGCACAATGCCGTAC GTTACGTACTTCCTT |
| R-NGS-RBSn | CATCCAGTCTATCAATTGTTGCCGGGAAGCTAGAGTAAGTAGTT CGCCAGTTAATAGTT |
| F-NGS-RBSm | CTTACTCTAGCTTCCCGGCAACAATTGATAGACTGGATGGAGGC GGATAAAGTTGCAGG |
| R-NGS-RBSm | TACGGCATTGTGCAAATGCCTAGATCGGAACAACACCGGTGAA CAGTTCTTCACCTATA |
| F-I | GCTTTCGATGGCGATGTCGCAGTCCAGA |
| R-I | TGCGACCAGCATGGTACGTGCCACGATA |
| F-II | ATCGGAGTTGGCGATGTCGCAGTCCAGA |
| R-II | ACTGAATCGCATGGTACGTGCCACGATA |
| F-III | TGTCGTACTGGCGATGTCGCAGTCCAGA |
| R-III | TCGCTGAAGCATGGTACGTGCCACGATA |
| F-IV | GAGCGTTTTGGCGATGTCGCAGTCCAGA |

| | |
|---|---|
| R-IV | CAGGTGCAGCATGGTACGTGCCACGATA |
| F-V | CGTTATGATGGCGATGTCGCAGTCCAGA |
| R-V | GACACAAGGCATGGTACGTGCCACGATA |
| F-*glcC* | TGCGCCTGCGCGTTGGTTCTCAACGCTCTCAATAAGCTTATGAAAGATGAACGTCGCCC |
| R-*glcC* | GGCATGTTGGTTTCCTACATTCAATTTTTTAGTCGCTTACTAACTCAGGTTCATCTCCA |
| F-pUC-AMP | ATTCGCTCGGCGGTGCCGCTGGAGATGAACCTGAGTTAGTAAGCGACTAAAAAATTGAA |
| R-pUC-AMP | AACAGAAAAATTGGTCCTACCTGTGCACGAGGTCCGGGAGAGTTTGTAGAAACGCAAAA |
| F-pglcD | TCCCGGACCTCGTGCACA |
| R- pglcD | GAGTAGGCTTCGCTTTGT |
| F-sfGFP | AATCAGCTGCCACACAACACAACAAAGCGAAGCCTACTCTTTAACTTTAAGAAGGAGAT |
| R-sfGFP | ATTCATTTCCCCAACGCGTCTGGCAAATCGTCGCCGCTATTCCCGAAGGCTATGGATCC |
| F-ffs | ATAGCCTTCGGGAATAGCGGCGA |
| R-ffs | AAGCTTATTGAGAGCGTTGAGAAC |
| F-cdaR-BJ08 | TCTAGAGAAAGACGAGATATACTAGATGGCTGGCTGGCATCTTGA |
| R-cdaR-BJ08 | CTAGTATATCTCGTCTTTCTCTAGAGCATAAGGAAGTACGTAACG |
| F-cdaR-Rc | TAAGCCGTGCATAACGGAGGACTTATGGCTGGCTGGCATCTTGAT |
| R-cdaR-Rc | AAGTCCTCCGTTATGCACGGCTTAGCATAAGGAAGTACGTAACGT |

| | |
|---|---|
| F-cdaR-Rg | ACAATTGGAGGAATAAGGTAAGCTTATGGCTGGCTGGCATCTTGA |
| R-cdaR-Rg | AAGCTTACCTTATTCCTCCAATTGTGCATAAGGAAGTACGTAACG |
| F-cdaR-RpR | GCCGTACGTTACGTACTTCCTTATGCTGTTTAACTTTAATAAGGAGATATAATGGCTGG |
| R-cdaR-RpR | CCAGCCATTATATCTCCTTATTAAAGTTAAACAGCATAAGGAAGTACGTAACGTACGGC |
| F-cdaR-RpT | AATTTCACACAGGAAACAGACCATGATGGCTGGCTGGCATCTTGA |
| R-cdaR-RpT | CATGGTCTGTTTCCTGTGTGAAATTGCATAAGGAAGTACGTAACG |
| F-cdaR-G10 | CGTACTTCCTTATGCTTTAACTTTAAGAAGGAGATATACATATGGCTGGCTGGC |
| R-cdaR-G10 | GCCAGCCAGCCATATGTATATCTCCTTCTTAAAGTTAAAGCATAAGGAAGTACG |
| R-gfp2-BJ04 | CTAGTCTGTCCCTTCTTTCTCTAGATGTTGCACTCCTGAAAATTC |
| R-gfp2-BJ08 | CTAGTATATCTCGTCTTTCTCTAGATGTTGCACTCCTGAAAATTCG |
| R-gfp2-Rc | AAGTCCTCCGTTATGCACGGCTTATGTTGCACTCCTGAAAATTCG |
| F-gfp2-Rg | ACAATTGGAGGAATAAGGTAAGCTTATGCGTATAGGTGAAGAACTG |
| R-gfp2-Rg | AAGCTTACCTTATTCCTCCAATTGTTGTTGCACTCCTGAAAATTCG |
| F-gfp2-RpR | TGTTTAACTTTAATAAGGAGATATAATGCGTATAGGTGAAGAACTGTTCACCGG |

| | |
|---|---|
| R-gfp2-RpR | TATATCTCCTTATTAAAGTTAAACATGTTGCACTCCTGAAAATTCGCGTTAGCC |
| R-gfp2-RpT | CATGGTCTGTTTCCTGTGTGAAATTTGTTGCACTCCTGAAAATTC |
| F-glc-BJ08 | TCTAGAGAAAGACGAGATATACTAGATGAAAGATGAACGTCGCCC |
| R-glc-ffs-BJ08 | CTAGTATATCTCGTCTTTCTCTAGAACCAACGCGCAGGCGCATTA |
| F-glc-Rc | TAAGCCGTGCATAACGGAGGACTTATGAAAGATGAACGTCGCCCT |
| R-glc-ffs-Rc | AAGTCCTCCGTTATGCACGGCTTAACCAACGCGCAGGCGCATTAT |
| F-glc-Rg | ACAATTGGAGGAATAAGGTAAGCTTATGAAAGATGAACGTCGCCC |
| R-glc-ffs-Rg | AAGCTTACCTTATTCCTCCAATTGTACCAACGCGCAGGCGCATTA |
| F-glc-ffs-RpR | GCGCGTTGGTTGTTTAACTTTAATAAGGAGATATAATGAAAGATGAACGTCGCCC |
| R-glc-ffs-RpR | TATATCTCCTTATTAAAGTTAAACAACCAACGCGCAGGCGCA |
| F-glc-RpT | AATTTCACACAGGAAACAGACCATGATGAAAGATGAACGTCGCCC |
| R-glc-ffs-RpT | CATGGTCTGTTTCCTGTGTGAAATTACCAACGCGCAGGCGCATTA |
| F-glc-M10 | TCTTAATCATGCGGAGGATCGTTTCATGAAAGATGAACGTCGCCC |
| R-glc-ffs-M10 | GAAACGATCCTCCGCATGATTAAGAACCAACGCGCAGGCGCATTA |
| F-glc-ffs-G10 | GCGCGTTGGTTTTAACTTTAAGAAGGAGATATACATATGAAAGATGAACGTCGCC |

| | |
|---|---|
| R-glc-ffs-G10 | ATGTATATCTCCTTCTTAAAGTTAAAACCAACGCGCAGGCGCATT |
| F-gfp-BJ04 | TCTAGAGAAAGAAGGGACAGACTAGATGCGTATAGGTGAAGAACT |
| R-gfp-BJ04 | CTAGTCTGTCCCTTCTTTCTCTAGAGAGTAGGCTTCGCTTTGTTG |
| F-gfp-BJ08 | TCTAGAGAAAGACGAGATATACTAGATGCGTATAGGTGAAGAACTG |
| R-gfp-BJ08 | CTAGTATATCTCGTCTTTCTCTAGAGAGTAGGCTTCGCTTTGTTGT |
| F-gfp-Rc | TAAGCCGTGCATAACGGAGGACTTATGCGTATAGGTGAAGAACTG |
| R-gfp-Rc | AAGTCCTCCGTTATGCACGGCTTAGAGTAGGCTTCGCTTTGTTGT |
| F-gfp-Rg | CACAATTGGAGGAATAAGGTAAGCTTATGCGTATAGGTGAAGAACTG |
| R-gfp-Rg | AAGCTTACCTTATTCCTCCAATTGTGAGTAGGCTTCGCTTTGTTGT |
| F-gfp-RpR | CTCTGTTTAACTTTAATAAGGAGATATAATGCGTATAGGTGAAGAACTGTTCACCGG |
| R-gfp-RpR | CCGGTGAACAGTTCTTCACCTATACGCATTATATCTCCTTATTAAAGTTAAACAGAG |
| F-gfp-RpT | AATTTCACACAGGAAACAGACCATGATGCGTATAGGTGAAGAACT |
| R-gfp-RpT | CATGGTCTGTTTCCTGTGTGAAATTGAGTAGGCTTCGCTTTGTTG |
| F-gfp-M10 | TCTTAATCATGCGGAGGATCGTTTCATGCGTATAGGTGAAGAACT |
| R-gfp-M10 | GAAACGATCCTCCGCATGATTAAGAGAGTAGGCTTCGCTTTGTTG |

| | |
|---|---|
| F-gfp-G10 | CTCTTTAACTTTAAGAAGGAGATATACATATGCGTATAGGTGAAGAACTGTTCACCGG |
| R-gfp-G10 | CGCATATGTATATCTCCTTCTTAAAGTTAAAGAGTAGGCTTCGCTTTGTTGTGTTGTG |
| F-Ac-gfp | ATGCGTATAGGTGAAGAAC |
| R-Ac-gfp | CTTTGTACAGTTCGTCCA |
| F-Ac-Para | CCATCCTGACGGATGGCCTTTTTGCGTTTCTACAAACTCATGCATGCCAGTTCTAGCAT |
| R-Ac-Para | CGGAACAACACCGGTGAACAGTTCTTCACCTATACGCATTTTTTATAACCTCCTTAGAG |
| F-Ac-Amp | CTGCTGGTATCACCCACGGTATGGACGAACTGTACAAAGTAAGCGACTAAAAAATTGAA |
| R-Ac-Amp | TAGCTCACTCATTAGGGTTATGCTAGAACTGGCATGCATGAGTTTGTAGAAACGCAAAA |
| yz-F-Ac | GACCTACGGTGTTCAGTG |
| yz-R-Ac | ACGACTTATCGCCACTGG |
| yz-F-glc | TCTGACCGGTAGCTAAAGAG |
| yz-R-glc | CTGAGAATAGTGTATGCGGC |
| R-gfp-ara-BJ04 | CTAGTCTGTCCCTTCTTTCTCTAGACGAATTCCCAAAAAAACGGG |
| R-gfp-ara-BJ08 | CTAGTATATCTCGTCTTTCTCTAGACGAATTCCCAAAAAAACGGG |
| F-ara-BJ08 | GCCATTCTAGAGAAAGACGAGATATACTAGGAAGAAACCAATTGTCC |
| R-ara-BJ08 | CTAGTATATCTCGTCTTTCTCTAGAATGGCTGAAGCGCAAAATGA |
| R-gfp-ara-Rc | AAGTCCTCCGTTATGCACGGCTTACGAATTCCCAAAAAAACGGGT |

| | |
|---|---|
| F-ara-Rc | TAAGCCGTGCATAACGGAGGACTTGAAGAAACCAATTGTCCATAT |
| R-ara-Rc | AAGTCCTCCGTTATGCACGGCTTAATGGCTGAAGCGCAAAATGAT |
| F-gfp-ara-Rg | GACAATTGGAGGAATAAGGTAAGCTTATGCGTATAGGTGAAGAAC |
| R-gfp-ara-Rg | AAGCTTACCTTATTCCTCCAATTGTCGAATTCCCAAAAAAACGGG |
| F-ara-Rg | GCCATACAATTGGAGGAATAAGGTAAGCTTGAAGAAACCAATTGTCC |
| R-ara-Rg | AAGCTTACCTTATTCCTCCAATTGTATGGCTGAAGCGCAAAATGA |
| F-gfp-ara-RpR | CGTGTTTAACTTTAATAAGGAGATATAATGCGTATAGGTGAAGAACTGTTCACCGG |
| R-gfp-ara-RpR | CTCCTTATTAAAGTTAAACACGAATTCCCAAAAAAACGGGTATGGAG |
| F-ara-RpR | GCGCTTCAGCCATTGTTTAACTTTAATAAGGAGATATAGAAGAAACCAATTGTCC |
| R-ara-RpR | TATATCTCCTTATTAAAGTTAAACAATGGCTGAAGCGCAAAATGATCCCCTGCTGC |
| R-gfp-ara-RpT | CATGGTCTGTTTCCTGTGTGAAATTCGAATTCCCAAAAAAACGGG |
| F-ara-RpT | AATTTCACACAGGAAACAGACCATGGAAGAAACCAATTGTCC |
| R-ara-RpT | CATGGTCTGTTTCCTGTGTGAAATTATGGCTGAAGCGCAAAATG |
| R-gfp-ara-M10 | GAAACGATCCTCCGCATGATTAAGACGAATTCCCAAAAAAACGGG |
| F-ara-M10 | TCTTAATCATGCGGAGGATCGTTTCGAAGAAACCAATTGTCCATA |

| | |
|---|---|
| R-ara-M10 | GAAACGATCCTCCGCATGATTAAGAATGGCTGAAGCGCAAAATGA |
| F-gfp-ara-G10 | CGTTTAACTTTAAGAAGGAGATATACATATGCGTATAGGTGAAGAACTGTTCACCGG |
| R-gfp-ara-G10 | CGCATATGTATATCTCCTTCTTAAAGTTAAACGAATTCCCAAAAAAACGGG |
| F-ara-G10 | GCGCTTCAGCCATTTTAACTTTAAGAAGGAGATATACATGAAGAAACC |
| R-ara-G10 | CATGTATATCTCCTTCTTAAAGTTAAAATGGCTGAAGCGCAAAATGATCCCCTGC |
| F-EL | CAGCCATCACCATCATCACCACAGCCAGGATCCGAATTCATGGCAGCTAAAGACGTAAA |
| R-EL | ACTTTCTGTTCGACTTAAGCATTATGCGGCCGCAAGCTTTTACATCATGCCGCCCATGC |
| F-ELzai | AAGCTTGCGGCCGCATAATGC |
| R-ELzai | GAATTCGGATCCTGGCTGTGG |
| F-ES | TAGTATATTAGTTAAGTATAAGAAGGAGATATACATATGATGAATATTCGTCCATTGCATGATCGCG |
| R-ES | TTTCGCAGCAGCGGTTTCTTTACCAGACTCGAGGGTACCTTACGCTTCAACAATTGCCAGAATGTCG |
| F-ESzai | AAGGTACCCTCGAGTCTGGTAAAGAAACC |
| R-ESzai | GGACGAATATTCATCATATGTATATCTCC |
| yz-F-ESL | GCAGCAGCCATCACCATCATC |
| yz-R-ESL | AGACCCGTTTAGAGGCCCCAA |
| F-RBSn81 | TAACCATGCATAAGGGAGCAGACTTATGGCTGGCTGGCATCTTGA |
| R-RBSn81 | AAGTCTGCTCCCTTATGCATGGTTAGCATAAGGAAGTACGTAACG |

| | |
|---|---|
| F-RBSm56 | CATGGGAAGGAAGGTTTCATGCGTATAGGT |
| R-RBSm56 | CCTTCCTTCCCATGATTAAGATGTTGCACT |
| F-RBSm97 | CATGAGGAGGCTGGTTTCATGCGTATAGGTG |
| R-RBSm97 | CCAGCCTCCTCATGATTAAGATGTTGCACT |
| F-RBSm117 | CATGTGAAGAATGGTTTCATGCGTATAGGT |
| R-RBSm117 | CCATTCTTCACATGATTAAGATGTTGCACTCC |

**Supplementary Table 6 DNA sequences**

| genes | Sequence (from 5' to 3') |
|---|---|
| The sgRNA of *lacZ* | TTAACTCGGCGTTTCATCTGTGG |
| *lacZ* upstream and downstream 500 bp from *E. coli* | GGCATCGTTCCCACTGCGATGCTGGTTGCCAACGATCAGATGG<br>CGCTGGGCGCAATGCGCGCCATTACCGAGTCCGGGCTGCGCGT<br>TGGTGCGGATATCTCGGTAGTGGGATACGACGATACCGAAGA<br>CAGCTCATGTTATATCCCGCCGTTAACCACCATCAAACAGGAT<br>TTTCGCCTGCTGGGGCAAACCAGCGTGGACCGCTTGCTGCAAC<br>TCTCTCAGGGCCAGGCGGTGAAGGGCAATCAGCTGTTGCCCGT<br>CTCACTGGTGAAAAGAAAAACCACCCTGGCGCCCAATACGCA<br>AACCGCCTCTCCCCGCGCGTTGGCCGATTCATTAATGCAGCTG<br>GCACGACAGGTTTCCCGACTGGAAAGCGGGCAGTGAGCGCAA<br>CGCAATTAATGTGAGTTAGCTCACTCATTAGGCACCCCAGGCT<br>TTACACTTTATGCTTCCGGCTCGTATGTTGTGTGAAATTGTGAG<br>CGGATAACAATTTCACACAGGAAACAGCTTAATAACCGGGCA<br>GGCCATGTCTGCCCGTATTTCGCGTAAGGAAATCCATTATGTA<br>CTATTTAAAAAACACAAACTTTTGGATGTTCGGTTTATTCTTTT<br>TCTTTTACTTTTTTATCATGGGAGCCTACTTCCCGTTTTTCCCGA<br>TTTGGCTACATGACATCAACCATATCAGCAAAAGTGATACGGG<br>TATTATTTTTGCCGCTATTTCTCTGTTCTCGCTATTATTCCAACC |

| | |
|---|---|
| | GCTGTTTGGTCTGCTTTCTGACAAACTCGGGCTGCGCAAATAC |
| | CTGCTGTGGATTATTACCGGCATGTTAGTGATGTTTGCGCCGTT |
| | CTTTATTTTTATCTTCGGGCCACTGTTACAATACAACATTTTAG |
| | TAGGATCGATTGTTGGTGGTATTTATCTAGGCTTTTGTTTTAAC |
| | GCCGGTGCGCCAGCAGTAGAGGCATTTATTGAGAAAGTCAGC |
| | CGTCGCAGTAATTTCGAATTTGGTCGCGCGCGGATGTTTGGCT |
| | GTGTTGG |
| BBa_J61104 | TCTAGAGAAAGAAGGGACAGACTAG |
| BBa_J61108 | TCTAGAGAAAGACGAGATATACTAG |
| $RBS_{cdaR}$ | TAAGCCGTGCATAACGGAGGACTT |
| $RBS_{glcC}$ | ACAATTGGAGGAATAAGGTAAGCTT |
| $RBS_{\mathrm{pRSF}}$ | TGTTTAACTTTAATAAGGAGATATA |
| $RBS_{\mathrm{pTrc99a}}$ | AATTTCACACAGGAAACAGACCATG |
| MCD10 | TCTTAATCATGCGGAGGATCGTTTC |
| g10RBS | TTTAACTTTAAGAAGGAGATATACAT |
| araBAD promoter | AAGAAACCAATTGTCCATATTGCATCAGACATTGCCGTCACTG |
| | CGTCTTTTACTGGCTCTTCTCGCTAACCAAACCGGTAACCCCGC |
| | TTATTAAAAGCATTCTGTAACAAAGCGGGACCAAAGCCATGAC |
| | AAAAACGCGTAACAAAAGTGTCTATAATCACGGCAGAAAAGT |
| | CCACATTGATTATTTGCACGGCGTCACACTTTGCTATGCCATAG |

| | |
|---|---|
| | CATTTTTATCCATAAGATTAGCGGATCCTACCTGACGCTTTTTA |
| | TCGCAACTCTCTACTGTTTCTCCAT |
| *araC* | TTATGACAACTTGACGGCTACATCATTCACTTTTTCTTCACAAC |
| | CGGCACGGAACTCGCTCGGGCTGGCCCCGGTGCATTTTTTAAA |
| | TACCCGCGAGAAATAGAGTTGATCGTCAAAACCAACATTGCG |
| | ACCGACGGTGGCGATAGGCATCCGGGTGGTGCTCAAAAGCAG |
| | CTTCGCCTGGCTGATACGTTGGTCCTCGCGCCAGCTTAAGACG |
| | CTAATCCCTAACTGCTGGCGGAAAAGATGTGACAGACGCGAC |
| | GGCGACAAGCAAACATGCTGTGCGACGCTGGCGATATCAAAA |
| | TTGCTGTCTGCCAGGTGATCGCTGATGTACTGACAAGCCTCGC |
| | GTACCCGATTATCCATCGGTGGATGGAGCGACTCGTTAATCGC |
| | TTCCATGCGCCGCAGTAACAATTGCTCAAGCAGATTTATCGCC |
| | AGCAGCTCCGAATAGCGCCCTTCCCCTTGCCCGGCGTTAATGA |
| | TTTGCCCAAACAGGTCGCTGAAATGCGGCTGGTGCGCTTCATC |
| | CGGGCGAAAGAACCCCGTATTGGCAAATATTGACGGCCAGTT |
| | AAGCCATTCATGCCAGTAGGCGCGCGGACGAAAGTAAACCCA |
| | CTGGTGATACCATTCGCGAGCCTCCGGATGACGACCGTAGTGA |
| | TGAATCTCTCCTGGCGGGAACAGCAAAATATCACCCGGTCGGC |
| | AAACAAATTCTCGTCCCTGATTTTTCACCACCCCCTGACCGCG |
| | AATGGTGAGATTGAGAATATAACCTTTCATTCCCAGCGGTCGG |
| | TCGATAAAAAAATCGAGATAACCGTTGGCCTCAATCGGCGTTA |

| | |
|---|---|
| | AACCCGCCACCAGATGGGCATTAAACGAGTATCCCGGCAGCA |
| | GGGGATCATTTTGCGCTTCAGCCAT |
| pglcD | TCCCGGACCTCGTGCACAGGTAGGACCAATTTTTCTGTTCTTAA |
| | CTCGCAAAACACGCACATCACGTAAGTGATTATGTTACATCAA |
| | TTTAACATTGAGTTAACCAAGACAAGGTCACAGAGCTGGAAA |
| | AAAAATGGTCTGACCGGTAGCTAAAGAGATAGACGAAAACGA |
| | AAAGCCCGCTTAATAACTGTTCACAGAAGCAGCGCGCAAAAA |
| | TCAGCTGCCACACAACACAACAAAGCGAAGCCTACTC |
| ffs promoter | ATAGCCTTCGGGAATAGCGGCGACGATTTGCCAGACGCGTTGG |
| | GGAAATGAATCTTCTTTTTCCATCTTTTCTTCCTGAGGTAATTT |
| | TTCAGCATAATCTGGAAAAACGCCCGAGTGAAGTCGCATTGCG |
| | CAAGAAACCAGCATCTGGCACGCGATGGGTTGCAATTAGCCG |
| | GGGCAGCAGTGATAATGCGCCTGCGCGTTGGTTCTCAACGCTC |
| | TCAAT |
| *glcC* | ATGAAAGATGAACGTCGCCCTATTTGCGAAGTGGTTGCAGAGA |
| | GTATCGAACGGTTAATTATCGACGGCGTACTGAAGGTCGGTCA |
| | GCCGCTTCCCTCGGAACGTCGACTGTGTGAAAAGCTCGGCTTC |
| | TCACGCTCCGCACTGCGTGAAGGGCTGACCGTGCTGCGCGGGC |
| | GCGGGATTATTGAAACGGCGCAGGGTCGCGATTCTCGTGTCGC |
| | ACGGCTTAATCGGGTGCAGGACACCAGCCCGCTGATCCATCTG |
| | TTCAGTACGCAGCCGCGAACGCTGTACGATCTGCTCGACGTTC |

| | |
|---|---|
| | GCGCATTACTGGAGGGCGAATCGGCAAGGCTGGCGGCAACGC |
| | TGGGAACGCAGGCTGATTTTGTTGTGATAACCCGCTGTTATGA |
| | AAAAATGCTCGCCGCCAGTGAGAACAACAAAGAGATTTCGCT |
| | GATCGAACATGCGCAGTTGGATCACGCTTTCCATCTCGCCATT |
| | TGTCAGGCTTCTCACAATCAGGTGCTGGTGTTTACGCTGCAAT |
| | CATTGACCGATCTGATGTTTAATTCAGTGTTTGCCAGCGTAAAT |
| | AATCTCTACCATCGACCACAGCAAAAAAAGCAGATCGATCGC |
| | CAGCATGCGCGGATCTACAACGCGGTGTTGCAGCGGCTGCCGC |
| | ACGTCGCCCAGCGCGCAGCACGCGATCATGTGCGGACCGTGA |
| | AAAAGAATCTCCACGATATCGAGCTGGAAGGCCACCATTTGAT |
| | TCGCTCGGCGGTGCCGCTGGAGATGAACCTGAGTTAG |
| *groEL* from *E. coli* | ATGGCAGCTAAAGACGTAAAATTCGGTAACGACGCTCGTGTGA |
| BL21 (DE3) | AAATGCTGCGCGGCGTAAACGTACTGGCAGATGCAGTGAAAGT |
| | TACCCTCGGTCCGAAAGGCCGTAACGTAGTTCTGGATAAATCTT |
| | TCGGTGCACCGACCATCACCAAAGATGGTGTTTCCGTTGCTCGT |
| | GAAATCGAACTGGAAGACAAGTTCGAAAATATGGGTGCGCAGA |
| | TGGTGAAAGAAGTTGCCTCCAAAGCGAACGACGCTGCAGGCG |
| | ACGGTACCACCACTGCAACCGTACTGGCTCAGGCTATCATCACT |
| | GAAGGTCTGAAAGCTGTTGCTGCGGGCATGAACCCGATGGACC |
| | TGAAACGTGGTATCGACAAAGCGGTTACCGCTGCAGTTGAAGA |
| | ACTGAAAGCGCTGTCCGTACCGTGCTCTGATTCTAAAGCGATTG |

CTCAGGTTGGTACCATCTCCGCTAACTCCGACGAAACCGTAGGT

AAACTGATCGCAGAAGCGATGGACAAAGTCGGTAAAGAAGGC

GTTATCACCGTTGAAGACGGTACCGGTCTGCAGGACGAACTGG

ACGTGGTTGAAGGTATGCAGTTCGACCGTGGCTACCTGTCTCCT

TACTTCATCAACAAGCCGGAAACTGGCGCAGTAGAACTGGAAA

GCCCGTTCATCCTGCTGGCTGACAAGAAAATCTCCAACATCCGC

GAAATGCTGCCGGTTCTGGAAGCTGTTGCAAAAGCAGGTAAAC

CGCTGCTGATCATCGCTGAAGATGTAGAAGGCGAAGCGCTGGC

AACTCTGGTTGTTAACACCATGCGTGGCATCGTGAAAGTCGCT

GCGGTTAAAGCACCGGGCTTCGGCGATCGTCGTAAAGCTATGC

TGCAGGATATCGCAACCCTGACTGGCGGTACCGTGATCTCTGAA

GAGATCGGTATGGAGCTGGAAAAAGCAACCCTGGAAGACCTG

GGTCAGGCTAAACGTGTTGTGATCAACAAAGACACCACCACTA

TCATCGATGGCGTGGGTGAAGAAGCTGCAATCCAGGGCCGTGT

TGCTCAGATCCGTCAGCAGATTGAAGAAGCAACTTCTGACTAC

GACCGTGAAAAACTGCAGGAACGCGTAGCGAAACTGGCAGGC

GGCGTTGCAGTTATCAAAGTAGGTGCTGCTACCGAAGTTGAAA

TGAAAGAGAAAAAAGCACGCGTTGAAGATGCCCTGCACGCGA

CCCGTGCAGCGGTAGAAGAGGGCGTGGTTGCTGGTGGTGGTGT

TGCGCTGATCCGCGTAGCGTCTAAACTGGCTGACCTGCGTGGTC

AGAACGAAGACCAGAACGTGGGTATCAAAGTTGCACTGCGTG

| | |
|---|---|
| | CAATGGAAGCTCCGCTGCGTCAGATCGTATTGAACTGCGGCGA<br>AGAACCGTCTGTTGTTGCTAACACCGTTAAAGGCGGCGACGGC<br>AACTACGGTTACAACGCAGCAACCGAAGAATACGGCAACATGA<br>TCGACATGGGTATCCTGGATCCAACCAAAGTAACTCGTTCTGCT<br>CTGCAGTACGCAGCTTCTGTGGCTGGCCTGATGATCACCACCG<br>AGTGCATGGTTACCGACCTGCCGAAAAACGATGCAGCTGACTT<br>AGGCGCTGCTGGCGGTATGGGCGGCATGGGTGGCATGGGCGGC<br>ATGATGTAA |
| *groES* from *E. coli* BL21 (DE3) | ATGAATATTCGTCCATTGCATGATCGCGTGATCGTCAAGCGTAA<br>AGAAGTTGAAACTAAATCTGCTGGCGGCATCGTTCTGACCGGC<br>TCTGCAGCGGCTAAATCCACCCGTGGCGAAGTGCTGGCTGTCG<br>GCAATGGCCGTATCCTTGAAAATGGCGAAGTGAAGCCGCTGGA<br>TGTGAAAGTTGGCGACATCGTTATTTTCAACGATGGCTACGGTG<br>TGAAATCTGAGAAGATCGACAATGAAGAAGTGTTGATCATGTC<br>CGAAAGCGACATTCTGGCAATTGTTGAAGCGTAA |
